# HBMAP: Bayesian inference of neural circuits from DNA barcoded projection mapping

**DOI:** 10.1101/2025.07.24.666115

**Authors:** Sara Wade, Edward Agboraw, Jinlu Liu, Huizi Zhang, Gülşen Sürmeli

## Abstract

Decoding how the brain routes information through its precise, long-range wiring remains a central challenge in neuroscience. Barcode-based mapping of axonal projections allows brain-wide, high-throughput investigation of projections at single-neuron resolution offering a powerful solution. However, principled methods for statistical analysis of barcode count data that can detect common projection rules and effectively integrate datasets across subjects are lacking. To address these issues, we developed a model-based clustering approach through hierarchical Bayesian mixtures which we call hierarchical Bayesian mapping of axonal projections (HBMAP). We show that the inferred model accurately reflects the features of the data and allows simultaneous identification of projection patterns and characterization of uncertainty that accounts for subject variability. Our study presents the first Bayesian approach to barcode-based projection mapping, offering a general solution for this class of data.

## Main

Mapping long-range connectivity is crucial to establish biological mechanisms of cognition and test hypotheses of brain dysfunction in disorders. Recently, mapping axonal projections of individual neurons, rather than bulk-labeling, has revealed unexpected diversity in projection patterns among otherwise similar neurons. These findings reshape cell type definitions by linking wiring patterns to molecular identity and positional coordinates [1–3]. Research in this area is facilitated by high-throughput techniques such as Multiplexed Analysis of Projections by Sequencing [MAPseq, 1], which reformulates projection mapping as a problem of sequencing rather than microscopy. With MAPseq, thousands of single-neurons are tagged in parallel with unique mRNA barcodes. RNA sequencing from tissue biopsies reveal complex branching and connectivity patterns across the brain by matching barcodes between soma and target areas where axon terminals are located. For each neuron, the barcode counts across the target regions indicate the relative strength of projection. Derivatives of MAPseq follow the same fundamental process but include innovations which expand their utility, for instance by incorporating in-situ transcriptomics [4–6]. These advances unlock an unprecedented ability to decode how genetic programs and circuit architecture jointly shape the functional roles of neurons within broader neural networks.

In MAPseq, the main goal is to identify robust *projection motifs* - recurring patterns of neuronal connectivity - from barcode count data. Two main analytical strategies have previously been used. The first transforms the data (e.g., normalizing and/or log-scaling barcode counts) followed by clustering using algorithms like k-means [7, 8] or k-medoids [9, 10], equipped with a chosen similarity measure, e.g. [11] use the cosine similarity and [12] use the Euclidean distance. This strategy ignores variability in projection strength and lacks interpretability of the clusters, making it hard to explain why clusters form, what they mean biologically and how preprocessing and data transformations affect results. Its heuristic nature prevents rigorous statistical inference, such as assessing cluster validity or how confidently each neuron belongs to a specific cluster. Although more flexible methods (e.g., density- or neural network–based clustering, [13–15]) can model complex cluster shapes, their black-box nature further reduces interpretability, which is necessary for meaningful biological insights.

The second strategy uses a binomial model [11, 16], which allows statistical testing for over- or underrepresented projection motifs compared to a null model, using one-sided binomial tests. Two versions of the null model exist: [16] assumes a uniform projection probability across all target regions and neurons, while [11] allows region-specific probabilities. Despite offering statistical rigor, this method also has drawbacks. It binarizes projection data, typically using thresholds of one or five barcodes, resulting in a clear loss of information (Fig. 1a-b). The choice of threshold and null model influences the findings, which can arbitrarily shift the categorization of motifs from significant to nonsignificant (demonstrated on data from [11] Fig. 1c-h). Moreover, the null model is effectively a statistical strawman, making it difficult to draw conclusions from rejection of the null hypothesis [17]; specifically, the significant motifs do not constitute the main ways in which neurons project. Indeed, they may contain a small number of neurons, while motifs with a large number of neurons are deemed insignificant and overlooked. For example, the over-represented motif in Fig. 1i reported in [11] has only two neurons, while the motif in Fig. 1j, is insignificant, despite containing many more neurons.

**Fig. 1:**
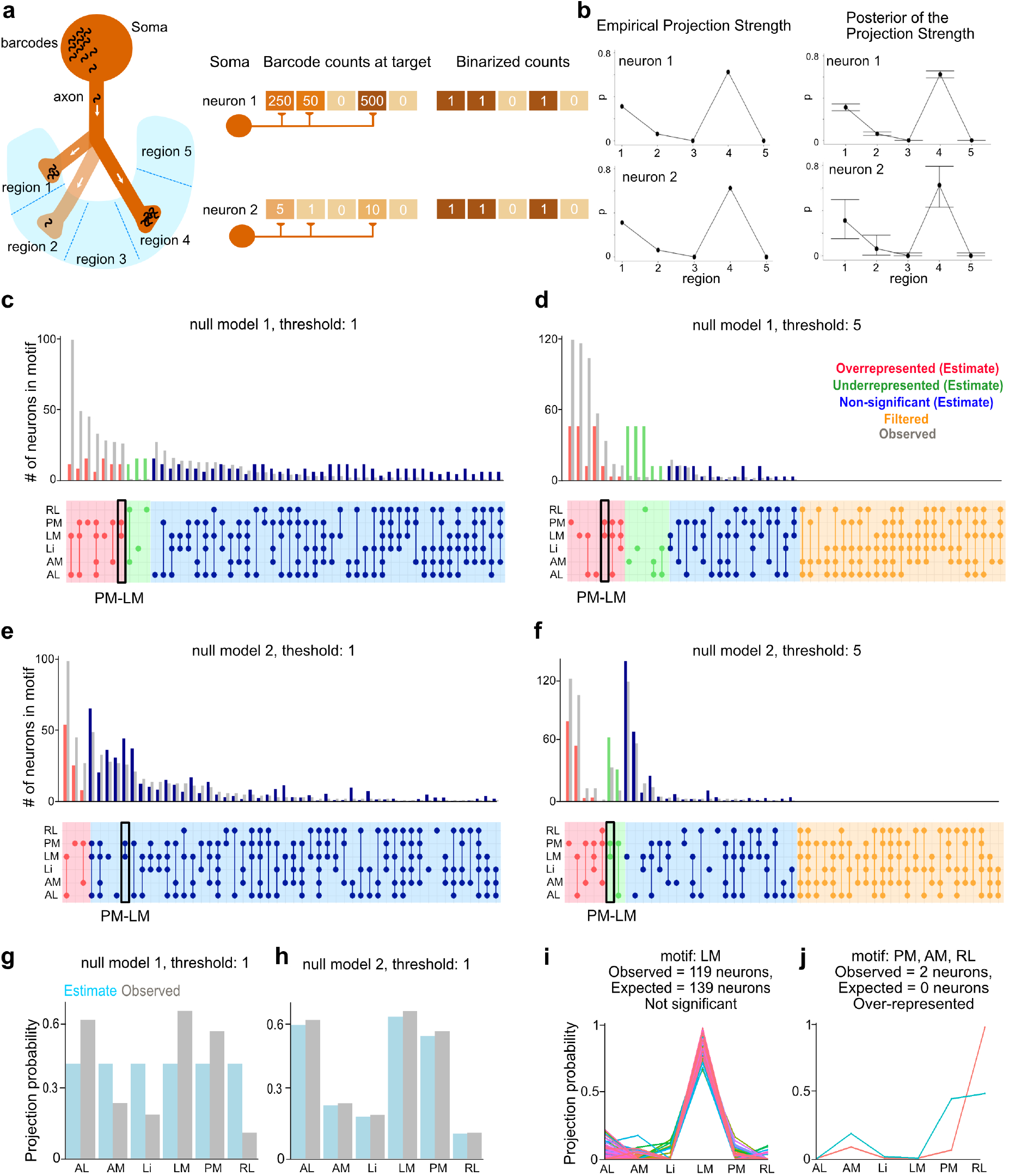
The choice of threshold and null model result in different categorizations of projection motifs in the binomial model. **a**. Schematics of the MAPseq barcode transport process from soma to the axon terminals (left). Binarization eliminates information on relative barcode counts across targets (right). **b**. Relative strength can be captured by the *empirical projection strength* (left), which divides counts by the total barcode count instead of simply binarizing. But, this fails to capture the uncertainty in a neuron’s projection strength arising from varying total count (neuron 1 high total count, neuron 2 low total count). Instead, the posterior distribution of the projection strength via hierarchical modeling (right) demonstrates increasing uncertainty (top to bottom) as the observed total counts decreases. **c-f**. Binomial model results for the primary visual cortex (V1) projections to the laterointermediate (LI), lateromedial (LM), anterolateral (AL), posteromedial (PM), anteromedial (AM), and rostrolateral (RL) areas [11]. Barplots compare the number of neurons observed in each projection motif (connected circles) against the expected number under the binomial model, for four different sets of parameters. Note the number and identity of over-represented motifs vary under the different thresholding schemes and null model assumptions in c-f (e.g. PM-LM). **g-h**. Barcharts comparing the observed and estimated per-region projection probabilities under two different binomial models, at a binarization threshold of 1. The first assumes a constant projection probability, while the second allows the projection probability to vary by region. **i-j**. Projection probability line charts of a small (*n* = 2 neurons) yet overrepresented and large (*n* = 119 neurons) yet insignificant motif identified in **f**, highlighting the disconnect between size and significance under the binomial model.

To overcome the limitations of these current methods—particularly their disregard for uncertainty and projection strength, and challenges in interpretability—we developed a probabilistic framework for projection motif analysis through hierarchical Bayesian mixture models (Hierarchical Bayesian mapping of axon projections, HBMAP). As opposed to simply pooling data across multiple brains, ignoring batch effects from experimental factors such as dissection inaccuracies and variation in virus injection site, the hierarchical Bayesian framework allows for data-driven borrowing of information across experiments. This offers a compromise between the two extremes of pooling the data and independent analyses of brains. HBMAP probabilistically clusters neurons, employing a Dirichlet-multinomial to directly model barcode counts within motifs and account for the overdispersion and sparsity present in the data. This avoids any transformation, accounts for variability, and improves interpretability. The inferred model accurately reflects features of real MAPseq data and allows us to simultaneously identify projection motifs, quantify uncertainty, measure differences between experiments, and generate synthetic datasets to investigate robustness and hypotheses. Thorough comparisons are provided on two datasets, highlighting the strength of HBMAP over existing approaches.

## Results

### Distinct features of MAPseq data pose analytical challenges

To illustrate the challenges inherent in modeling MAPseq data, we consider the dataset from [11] characterizing single-neuron outputs from the primary visual cortex (V1) to various targets in secondary visual domains. An exploratory analysis highlights challenges in modeling the data. The empirical projection strengths, the neuron’s barcode counts divided by its total count across regions, show multimodality and high dispersion (Fig. 2a), along with sparsity (degeneracy to the edges of the simplex) in the scatter plot (Fig. 2b). Additionally, some differences are evident across brains (Fig. 2c). These properties violate the assumptions of many standard statistical or clustering techniques (e.g., unimodality, equidispersion, pooling, estimates on the boundary of parameter spaces).

**Fig. 2:**
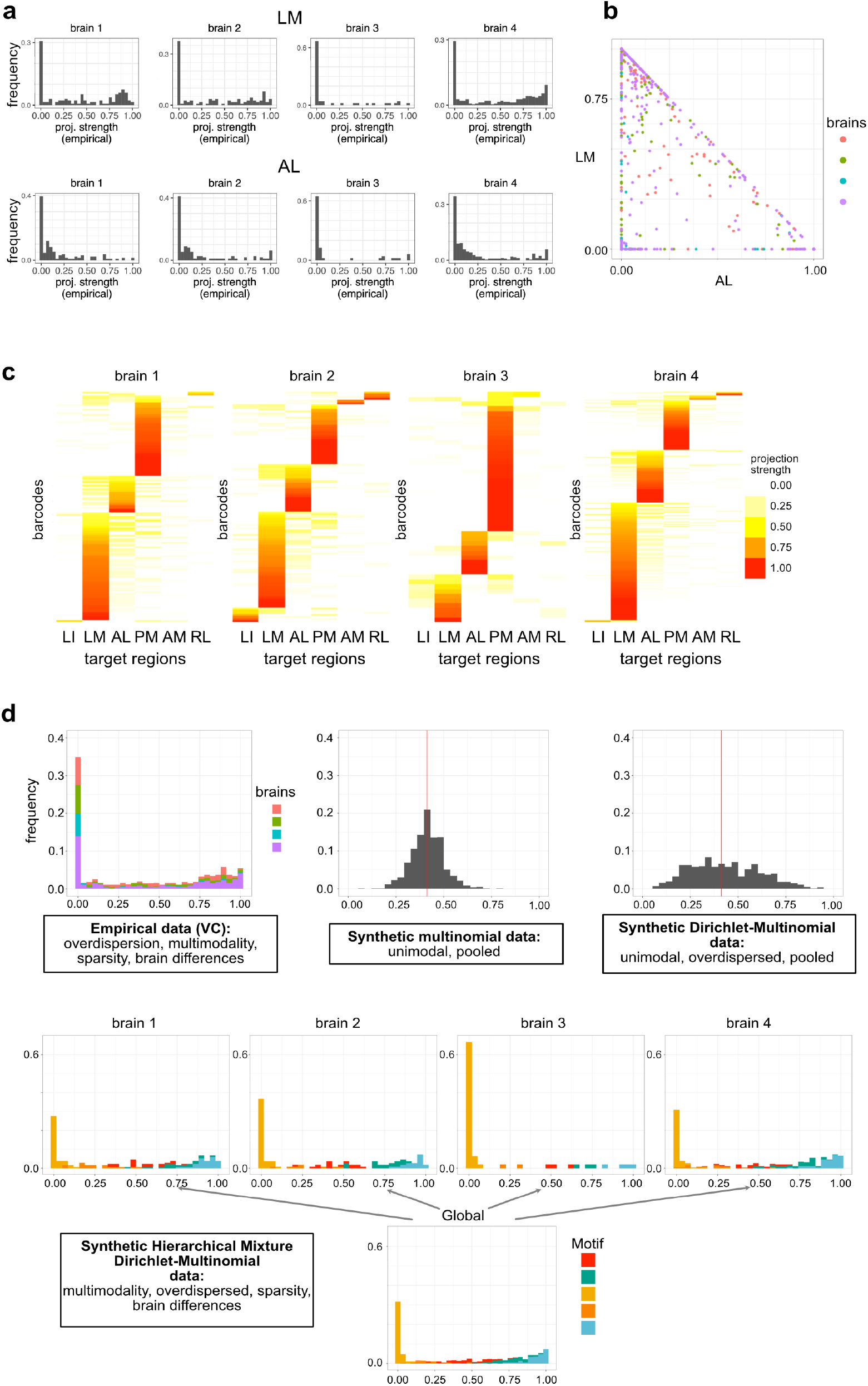
Barcode count data are sparse, overdispersed, multimodal, and variable across brains, making standard multinomial or Dirichlet-multinomial models inappropriate. **a**. Histograms of the empirical projection strengths (barcode counts divided by the total count) of V1 to LM and AL in all brains used in [11]. These plots indicate the presence of across sample, brain-level differences, multimodality and overdispersion in the data. **b**. Scatter plot of empirical projection strengths to AL (x-axis) and LM (y-axis), illustrating sparsity in the data. **c**. Heatmaps visualizing the empirical projection strengths of each barcode. Notable brain-level differences in projection are observed. **d**. Histograms comparing the distribution of empirical projection strengths to the LM region against those generated by a multinomial, Dirichlet-multinomial, and hierarchical Dirichlet-multinomial mixture model fitted to the data from [11]. Unlike other models, the hierarchical mixture model captures the multimodality, overdispersion, sparsity and brain-level differences seen in the empirical data.

### Overview of HBMAP

The challenging features of MAPseq data make it essential to adopt models that can explicitly account for its distinct structure. By building a custom model, our goal is to characterize the estimated projection strengths of individual neurons across brain regions, while rigorously accounting for uncertainty in these estimates. For each neuron, barcode counts are observed, denoted by **y**_*i,·,m*_ = (*y*_*i*,1,*m*_, …, *y*_*i,R,m*_) (for neuron *i* = 1, …, *n*_*m*_ in brain *m* = 1, …, *M*, across the *R* target regions). These counts reflect the neuron’s unknown projection strength vector 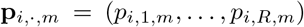. Each element *p*_*i,r,m*_ measures the probability of projecting to each target brain area relative to all other target areas considered, as such *p*_*i,r,m*_ ≥ 0 and *p*_*i*,1,*m*_ + … *p*_*i,R,m*_ = 1.

While empirical proportions provide a naive estimate of projection strength, they fail to quantify confidence, especially when barcode counts are sparse. Specifically, the empirical projection strengths are computed as the proportions of barcode counts 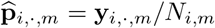 (where 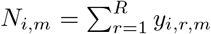is the neu-ron’s total barcode count), but this yields only point estimates with no notion of confidence or uncertainty. Instead, HBMAP models the barcode counts directly to account for uncertainty in the unobserved projection strengths (illustrated in Fig. 1b).

HBMAP models the observed barcode counts as outcomes of a multinomial process, which reflects the probabilistic nature of barcode targeting under uncertainty. Each neuron’s barcode counts are conceptualized as a series of *N*_*i,m*_ independent Bernoulli experiments (total number of barcodes that terminate in the target areas), where in each experiment a barcode is transported along branches to axon terminals in the *r*th target region with probability *p*_*i,r,m*_ (illustrated in Fig. 1a). Thus, the barcode counts follow a multinomial distribution,

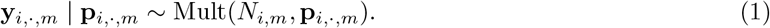

However, as the projection strength **p**_*i*,·,*m*_ in (1) is unknown, additional assumptions or hierarchical layers to the model are needed.

### Projection variability is better modeled by hierarchical than multinomial frameworks

Assuming all neurons share the same projection pattern leads to a simple multinomial model, but this approach is too restrictive and fails to capture biological diversity. Specifically, if all neurons have the same projection strength (**p**_*i,·,m*_ = **q** for all neurons) in (1), this implies that

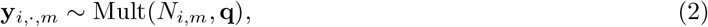

where **q** = (*q*_1_, …, *q*_*R*_) belongs to the simplex of dimension *R −* 1. In this model, the empirical projection strengths would be centered at **q**, i.e. 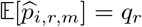for all neurons, with variance 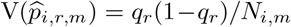. This multinomial model is overly simplistic, as illustrated by Fig. 2d, which highlights the disagreement between observed empirical projection strengths for the data from [11] and those obtained for synthetic data from this simple multinomial model fitted to this data.

To accommodate neuron-level variation, the Dirichlet-multinomial model adds a hierarchical layer that introduces overdispersion around a shared population mean. The Dirichlet-multinomial generalizes the multinomial (2), assuming:

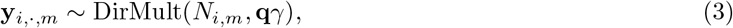

with an additional parameter *γ* > 0. The Dirichlet-multinomial is a compound distribution, which can be expressed as:
xs

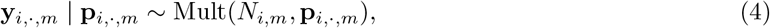

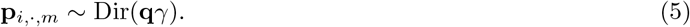

Thus, it includes a hierarchical layer to model the unknown projection strengths and naturally assumes that not all neurons share the same projection strength but have slight differences, for example, due to false positives and negatives that may arise in data collection, although at relatively low rates [11]. Compared to the multinomial model (2), the empirical projection strengths are still centered at **q**, i.e. 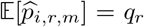, but the variance is increased: 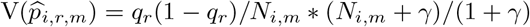. This allows overdispersion, with smaller *γ* producing higher dispersion in the counts, and the multinomial model (2) obtained in the limiting case *γ* → ∞. Indeed, the fitted Dirichlet-multinomial model to the data from [11] (Fig. 2d) better captures the overdispersion present in the data.

Moreover, to account for variability associated to the observed barcode counts, we need a formulation that can express uncertainty in the projection strengths, reflecting greater confidence when barcode counts are high. This is naturally incorporated in the hierarchical formulation in (4)-(5), which allows a Bayesian interpretation, where the *posterior* of the neuron’s projection strength given the observed counts is:

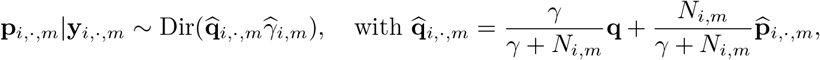

and 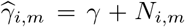. The neuron’s posterior expected projection strength, 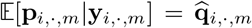, is a weighted average between the empirical projection strength and the prior global mean **q**, with posterior variability 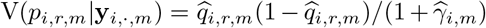. When the total barcode count *N*_*i,m*_ is small, there is less data and the posterior is more influenced by the prior with higher variability. Instead, for large total barcode count, the data dominate, and the posterior becomes closely aligned with the empirical projection strength with low variability. Thus, the Dirichlet-multinomial model (3) accounts for uncertainty in the neuron’s projection strength due to the total number of counts (illustrated in the right of Fig. 1b, with higher total count and less uncertainty for the first neuron compared to the second).

Despite these advantages, we again observe that Dirichlet-multinomial model (3) is too simple and overdispersion is not enough (Fig. 2d). In particular, the unimodal nature of the Dirichlet-multinomial is unable to account for the multiple modes of projection (different peaks in the histogram of the observed data in Fig. 2d).

### Mixtures of Dirichlet-multinomials capture the unique features of the data

To capture the multimodality evident in the data and flexibly model the distribution of neural projections, HBMAP employs mixtures of Dirichlet-multinomials (Fig. 3), which can be hierarchically expressed as:

**Fig. 3:**
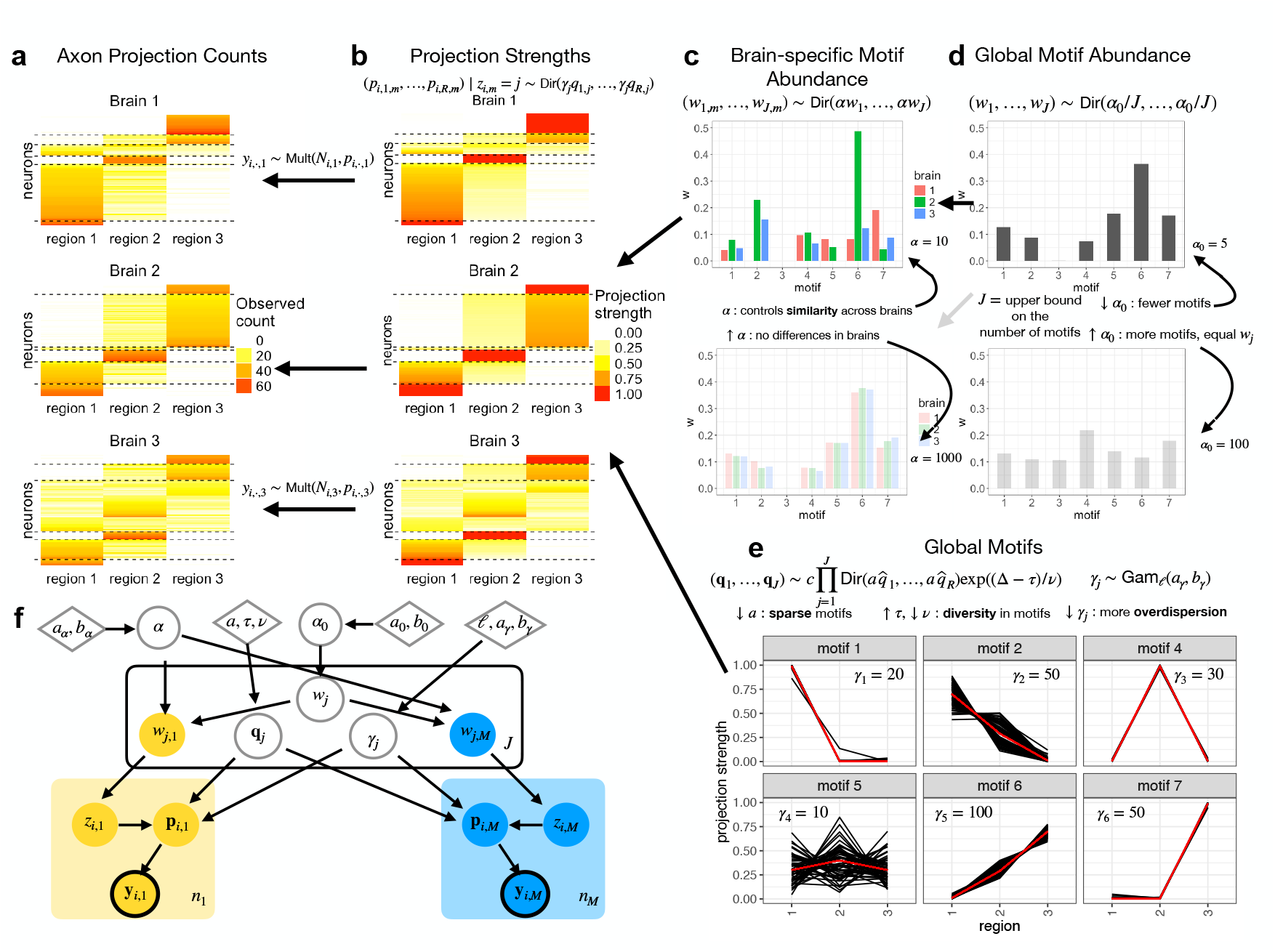
The generative flow of HBMAP, from the motifs and global weights on the right to brain-specific weights, projection strengths, barcode counts on the left. **a-b**. Heatmaps of the generated barcode counts in **a** and projection strengths in **b** for synthetic data generated by HBMAP, with dashed lines separating the clusters. The hierarchical framework assumes the observed barcode counts follow a multinomial model, depending on the projection strengths, which are in turn produced from the underlying mixture components, characterized by the brain-specific weights (**c**) and projection motif parameters (**e**). **c-d**. Barcharts of the brain-specific and global motif abundances in the synthetic data. The brain-specific motif abundances are centered around the global abundances, with the parameter *α* controlling variability across brains (larger values of *α* imply more similarity and pooling of information across brains) and the parameter *α*_0_ controlling the number of motifs. **e**. Projection strength line charts for the six global motifs observed. The global motifs are characterized by the mean projection strength **q**_*j*_ (in red) and dispersion parameter *γ*_*j*_; the **q**_*j*_ follow a repulsive Dirichlet model (smaller values of the hyperparameter *a* encourage sparser motifs, while larger *τ* and smaller *v* increase repulsion and diversity in **q**_*j*_ across motifs). Small values of *γ*_*j*_ (e.g. *γ*_*j*_ = 10 for cluster 4) corresponding to more variability in the projection strengths within the motif, and large values (e.g. *γ*_*j*_ = 50 for cluster 6) correspond to small variability. **f**. Graphical model of HBMAP. Observed neuron counts are outlined in black circles. Circles with white fills indicate global parameters and colored fills indicate brain-specific parameters. Diamonds denote fixed hyperparameters.

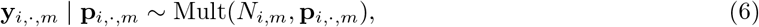

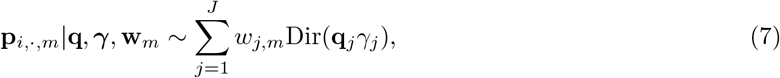

where each component *j* refers to one of the distinct clusters (motifs) of neuron projection patterns, with **q**_*j*_ = (*q*_1,*j*_, …, *q*_*R,j*_) representing the mean projection strength of the *j*th motif; *γ*_*j*_ > 0 controlling the variability of the projection strengths within each motif; and *w*_*j,m*_ reflecting the proportion of neurons in brain *m* that belong to the *j*th motif. Let **q** = (**q**_1_, …, **q**_*J*_), ***γ*** = (*γ*_1_, …, *γ*_*J*_), and **w**_*m*_ = (*w*_1,*m*_, …, *w*_*J,m*_) collect the parameters across all components. In this case, mixtures of Dirichlet-multinomials allow for both neuron-specific projection strengths (e.g. due to small differences from false positives and negatives) and multiple types of projection motifs across neurons. Fig. 2d compares the observed data from [11] to the fitted hierarchical mixture of Dirichlet-multinomials, highlighting the abilities of HBMAP to capture the unique features of neural projections.

### A hierarchical framework to integrate data across brains

The integration of data across brains is crucial for sound statistical analysis of projection motifs and their reliability. Existing approaches simply pool the data, which ignores across sample variability and inflates sample sizes leading to overconfident conclusions. In fact, brain-level differences are anticipated, specifically, in the proportion of neurons in each projection motif, for various reasons including minor variation in the injection site or inconsistencies in brain dissections. HBMAP accounts for this by allowing this proportion, called the motif abundance **w**_*m*_ = (*w*_1,*m*_, …, *w*_*J,m*_), to vary across brains, yet the projection patterns defining the motifs are shared by having common parameters (**q**_*j*_, *γ*_*j*_).

The hierarchical framework of HBMAP (illustrated in Fig. 3) is a compromise between the two extremes of pooling data and independent analyses across brains. It allows for data-driven borrowing of information across brains, by assuming each **w**_*m*_ is generated hierarchically from a Dirichlet distribution centered at the *global motif abundances* **w** = (*w*_1_, …, *w*_*J*_):

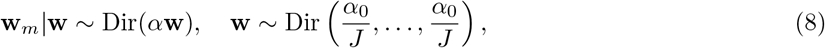

where *α* ∈ ℝ^+^ and *α*_0_ ∈ ℝ^+^ control variability in the motif abundances across brains and the total number of motifs, respectively. Fig. 3c illustrates the role of *α*, where variation in motif abundance decreases from top to bottom as *α* increases, with no brain differences and pooling in the limit as *α* → ∞. Fig. 3d demonstrates the effect of *α*_0_; HBMAP belongs to class of sparse overfitted mixtures [18–21], which infer the unknown number of motifs through a sparsity-promoting Dirichlet prior on the weights **w** with parameter *α*_0_. Here, *J* represents an upper bound on the number of motifs, and as *α*_0_ increases from top to bottom in Fig. 3d, the weights become less sparse and the number of motifs increases. HBMAP also infers these parameters (see Methods), providing statistical tools to both assess the number of motifs and quantify experimental variability.

### Applications of the HBMAP Toolkit: Comprehensive framework for motif analysis, uncertainty estimation, data integration, and synthetic data generation

HBMAP offers a toolkit to infer projection motif structure, estimate confidence in assignments of neurons to identified clusters, quantify experimental variability, and generate synthetic data. We demonstrate these strengths on two datasets [11, 12] by comparing HBMAP’s clustering to other methods, evaluating robustness, and showcasing tools for interpreting results and uncertainties.

#### Probabilistic labeling of projection motifs based on abundance and confidence

To interpret each cluster biologically, we assess how confidently we can determine its characteristic projection motif. HBMAP provides estimates of the mean projection strength for each motif along with credible intervals (CIs) (Supplementary Fig. 1). As the mean projection strength will not be exactly zero, HBMAP uses a probabilistic rule, where a region is considered a target only if the projection strength of neurons within the motif exceeds a threshold with sufficiently high probability (plot labels in Supplementary Fig. 1). Therefore, unlike other clustering methods, HBMAP provides biologically interpretable, uncertainty-aware projection motif labels.

### HBMAP overcomes sensitivities of heuristic clustering to preprocessing choices

The definition of projection motifs in heuristic clustering methods is sensitive to preprocessing choices and similarity metrics, often leading to varied biological interpretations. Fig. 4a illustrates this by comparing a subset of the motifs found by k-medoids (cosine similarity with max-scaling) and k-means (Euclidean distance with log-normal scaling), both set to identify 8 clusters as in [11]. For example, in k-means, the third most abundant cluster consists of neurons projecting primarily to AL, but k-medoids combines these neurons with those that also have weak projections to LM, losing the distinct AL cluster. This demonstrates a general challenge in choosing transformations and similarity measures in heuristic clustering algorithms due to the lack of interpretability in how they define a cluster. HBMAP does not require such choices. It uses a Dirichlet-multinomial distribution for each projection motif, which, as previously motivated, is an intuitive model of barcode counts.

**Fig. 4:**
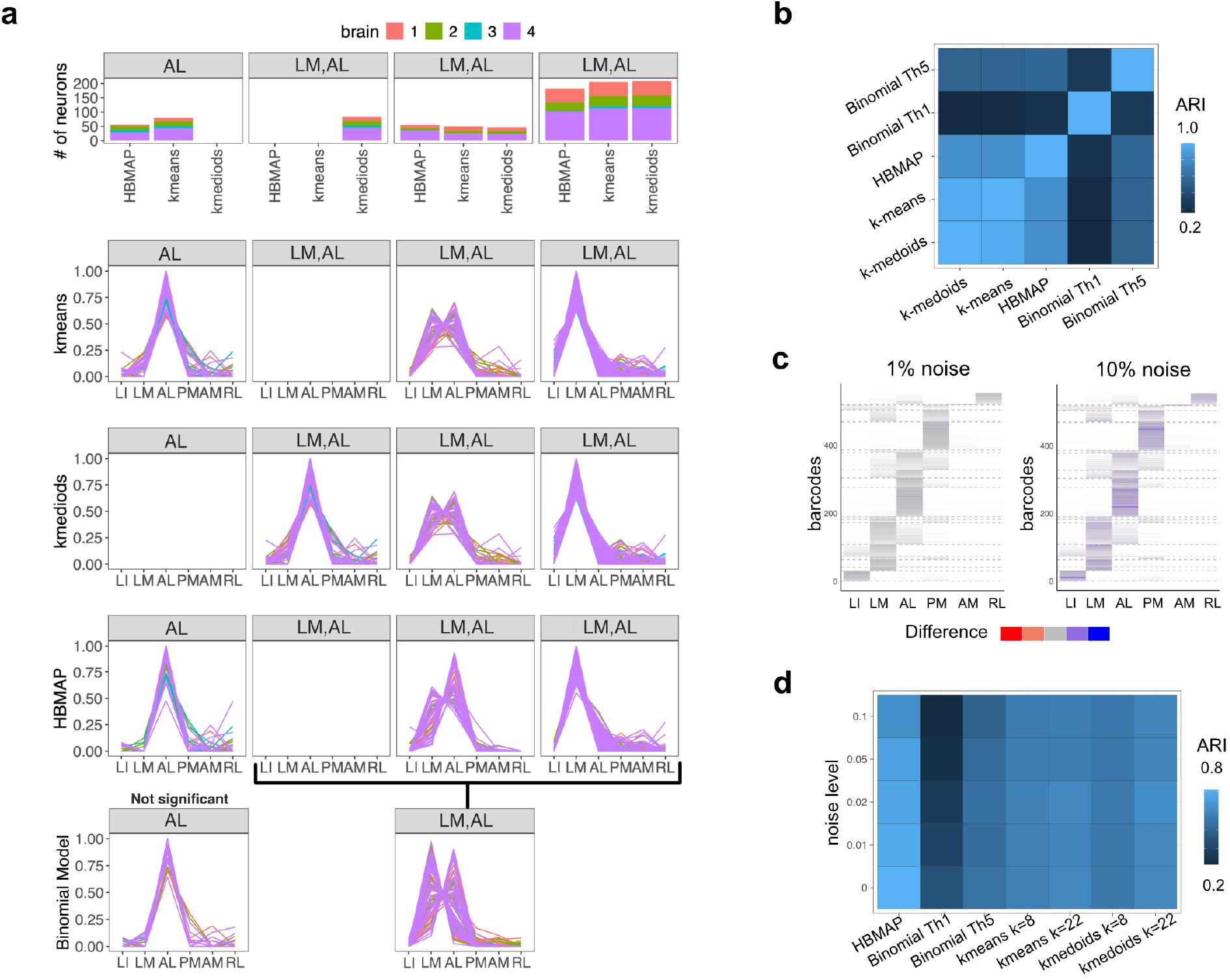
HBMAP versus alternative motif analysis methods. **a**. Comparison of a subset of the motifs (columns) identified by k-means, k-medoids, HBMAP, and the binomial model for the data from [11] (additional motifs for k-means, k-mediods, and HBMAP are shown in Supplementary Fig. 2. The binomial model uses the same choice of null model and threshold as in [11], with all motifs in Fig. 1f.). Top row: number of neurons in each motif across the three clustering methods (k-means, k-medoids, HBMAP). Bottom rows: empirical projection strength of neurons within each motif (column) for each model (row). For k-means and k-medoids, the target regions of each motif (listed in the panel titles, along with the cluster number) are determined by thresholding the average empirical projection strength across neurons in the motif at 0.05. For HBMAP, the regions of each motif are identified by a probabilistic rule, ensuring that neurons in the motif project to the region with sufficiently high probability. **b**. Adjusted Rand Index (ARI) illustrating the overlap in the clustering solutions found by the binomial, k-means, k-medoids and HBMAP models, with a value of 1 indicating equivalent clustering solutions and values close to zero when the two clusterings have many disagreements. **c**. Heatmaps illustrate the impact of dissection noise on synthetic data generated by HBMAP. The generative nature of HBMAP permits creation of realistic synthetic data, with known motif structure. To investigate sensitivity of the models, dissection noise was added to the synthetic data by randomly allocating barcode counts to alternate regions. Empirical projection strength is indicated by opacity. Differences in projection strength between the noise condition and the original data are indicated by color. Dashed lines identify the true clustering. **d**. ARI between the ground truth and inferred motif structure by different models, illustrating the impact of simulated dissection noise.

The main motifs discovered by HBMAP, closely match those from k-means and k-medoids (Fig. 4a and Supplementary Fig. 2). Unlike the binomial model which disregards nuances in projection strength, all three approaches reveal two sub-motifs to (LM, AL), with a large motif that has higher projection strength to LM and lower to AL (↑LM, ↓AL) and a smaller motif showing the inverse (↓LM, ↑AL). However, there are differences in the strength of projection for the smaller (↓LM, ↑AL) motif, which is higher to AL relative to LM in HBMAP. In addition, HBMAP finds a (AL) motif, which is present in all four mice and absent from k-medoids and classified as insignificant in the binomial model (Fig. 1f).

### Generating synthetic neural projection data with HBMAP to evaluate model fit and robustness

The generative capacity of HBMAP allows the simulation of realistic synthetic data (drawn from the posterior predictive distribution), which is useful to assess model fit, using posterior predictive checks [22], and to test robustness or hypotheses against a setting with known ground truth. For the former, posterior predictive checks demonstrate good fit of HBMAP to the observed data (Supplementary Fig. 3), and for the latter, we consider an illustration to study the effect of noise from dissection inaccuracies.

To mimic dissection noise in the synthetic data, a small percentage of barcode counts is randomly reallocated to the other regions (Fig. 4c). As the noise level increases, there are fewer neurons with exactly zero barcode counts in specific regions. The binomial model with a threshold of 1 is the most affected, with almost no agreement to the true clustering (Adjusted Rand Index (ARI) close to 0 in Fig. 4d). Although increasing the threshold to 5 provides an improvement, the binomial model still has the lowest agreement with the true clustering. For k-means and k-medoids, we consider 8 and 22 clusters (corresponding to the number of clusters in [11] and the true number in the synthetic data, respectively). These methods are more robust to dissection noise; however HBMAP better recovers the true clustering at all noise levels.

### HBMAP’s probabilistic framework enables rigorous projection motif analysis with uncertainty quantification

HBMAP enables reliable conclusions from MAPseq data by quantifying confidence in projection motifs, a critical step ignored by previous methods. Where others provide only a single clustering solution, HBMAP uses a probabilistic framework to account for variability across samples and individuals. To distinguish robust organizational principles from statistical noise, HBMAP identifies *prominent* motifs that are consistent features of the broader cell population, beyond just the current dataset. HBMAP calculates the posterior probability that the motif’s overall abundance (*w*_*j*_) across the entire population is greater than a set threshold (*ϵ*). For data investigating projections of V1 (with *ϵ* = 0.02), six motifs are classified as prominent (Fig. 5a,b). We therefore refine the clustering solution and propose that (AL), (PM), (↑LM, ↓AL), (↓LM, ↑AL), (↑LM, ↓AL, PM) and (LI, ↑LM, PM) best represent the projection patterns emerging from V1. Neurons targeting either AL or PM are abundant across all samples. This observation reinforces the conclusion from [11] that these areas receive distinct functional inputs from V1. Whereas they reached this conclusion by identifying the dual-projection PM-AL motif as underrepresented, our findings highlight the prominence of specialized, unicasting projections from V1 to each area individually. While [11] identified the PM-AM motif as overrepresented, our analysis reveals that PM and AM are instead embedded within broader, more diverse projection motifs that often include AL and LM (Fig. 5a). Although less abundant, these clusters highlight the capacity of V1 neurons to broadcast widely across the cortex. Consistent with [11], we observe motifs that include projections to a mixture of targets from the dorsal (AL, PM) and ventral (LM, LI) visual streams in Fig. 5a. However, we suggest that one of the predominant trifurcation motifs involves projections to LM, AL, and PM which was not discovered by the binomial analysis presented in [11]. Despite differences in specific motif composition, our findings support the conclusion that V1 neurons can make targeted as well as distributed projections, indicating a flexible routing architecture.

**Fig. 5:**
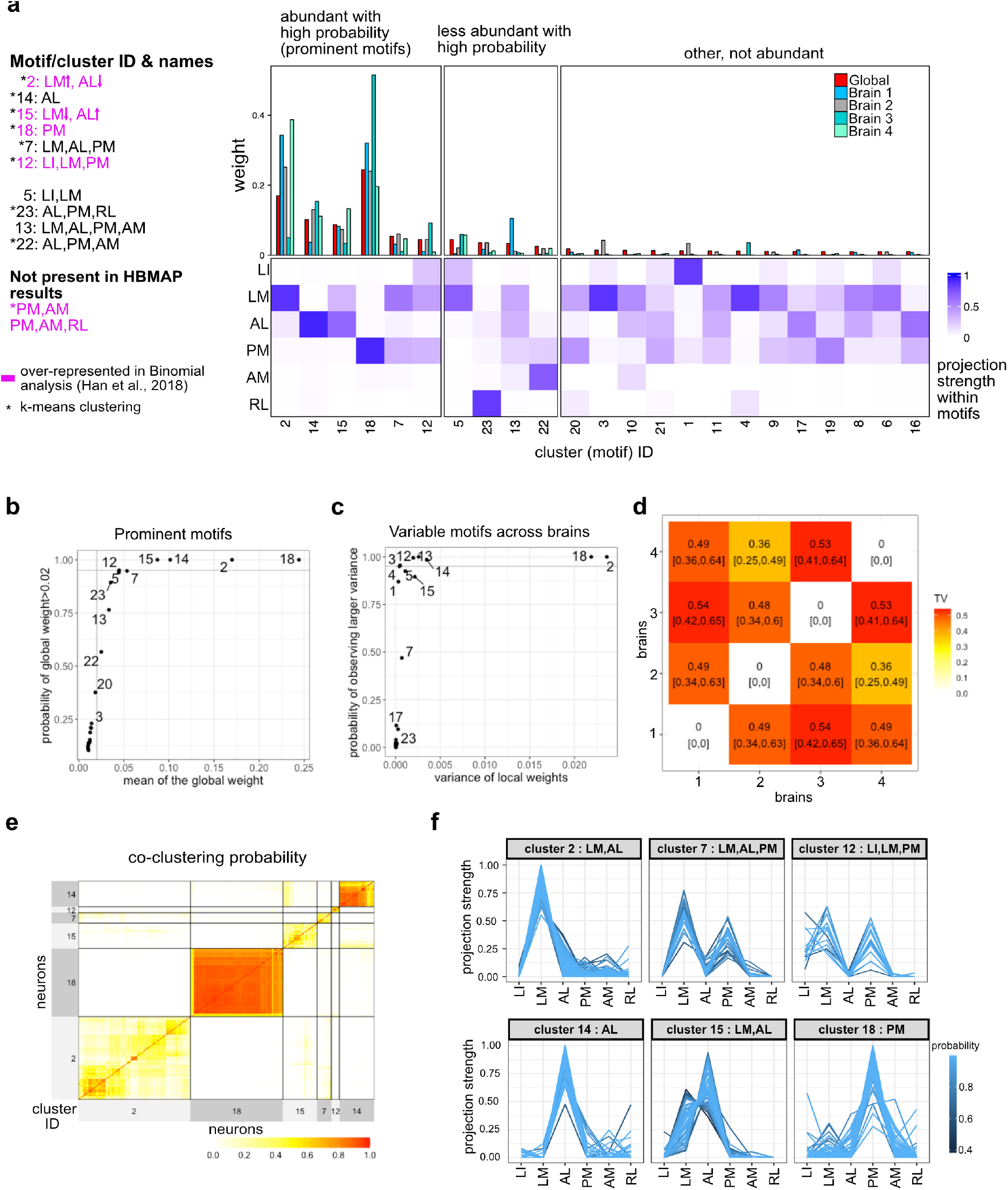
HBMAP estimates uncertainty in the motifs and assesses inter-individual variability. **a**. A list of projection motifs with high probability discovered by HBMAP (left). Heatmap of the estimated projection strengths to six target brain regions for each motif. Color intensity (blue to white) reflects relative projection strength to target areas. The estimated global and brain-specific weight attached to each motif are visualized as bar charts above the heatmaps. Motifs are grouped into prominent (global weight > 0.02, high probability), intermediate (global weight > 0.02, low probability), and other motifs. **b-c**. Scatter plots identifying prominent (left) and variable (right) motifs, with numbers indicating motif IDs. **b**. Estimated global motif weight vs. the posterior probability of the weight exceeding 0.02. **c**. Estimated variance of motif weights across individuals vs. the probability that this variance is greater than expected under a null model (no inter-individual difference). **d**. Total variation (TV) distance, illustrating the extent to which the inferred motif structures diverge across samples, with the extreme values of 0 and 1 indicating complete or no overlap in motif structure respectively. **e-f**. Illustration of uncertainty in motif structure through the posterior similiarity matrix (**e**) and assignment probabilities (**f**). **e**. Posterior similarity matrix of neurons in the prominent motifs, representing the probability of co-clustering. Lines separate neurons into motifs, and warmer colors indicate a higher probability of two neurons co-clustering. **f**. Projection strength profiles for neurons within prominent motifs. Lines are colored by the neuron’s posterior probability of assignment to that motif.

#### Quantifying inter-animal variability in projection motif structure

As opposed to pooling data across brains, HBMAP accounts for inter-individual variability through the hierarchical framework, allowing for new tools to assess this variability. We focus on two: 1) identifying variable motifs, whose abundance varies across brains, and 2) quantifying the overall similarity in the motif structure across brains.

First, to assess motifs with varying abundance across brains, for each motif, HBMAP computes the posterior expected variance of the local abundance *w*_*j,m*_ across brains. This is compared to a null setting, which assumes no differences across brains and that the neurons’ motif assignments are generated identically based on the global motif abundance. Variability of projection motifs are evaluated by computing the probability that the variance given the data is larger than the variance under the null setting (Fig. 5c). Two motifs stand out with their high variance: cluster 18 (PM) and cluster 2 (↑LM, ↓AL). The high variance is mostly due to brain 3, which presents a starkly inverted phenotype, where the PM motif dominates and the (↑LM, ↓AL) motif is suppressed, a pattern directly contrary to all other samples (Fig. 5a, bar plots). Several other motifs, including one associated with the ventral processing stream (↓LI, ↑LM), were abundant, with probability close to the 0.95 threshold, yet failed to meet the statistical threshold for prominence due to its complete absence in brain 1. Therefore, the conclusion about the absence of specific projection motifs, dedicated to the ventral processing stream in [11] could be revisited. Our analysis suggests that such a motif is indeed present, but its inconsistent expression across a limited cohort can cause it to fall below the threshold for statistical significance. This is either an indication of the stochastic nature of this projection motif, or experimental factors for instance variability amongst injection sites and types of infected neurons. To quantify the overall difference in the organization of the circuit between brains, we performed a pairwise comparison of the complete structures of the inferred motif using the total variation distance (TV) [23, Chapter 1]. TV scores (Fig. 5d) provide a metric to assess population-wide circuit variability and objectively identify biological outliers. Through the Bayesian framework, we inferred the expected TV distance, along with CIs, between any pair of brains and found further support that brain 3 is the most different from the other brains, although due to the small number of neurons (*n*_3_ = 48), the CIs are wide and overlap with the other pairs of brains included in the comparison.

#### Clustering uncertainty

Standard clustering algorithms often provide an optimal result, forcing messy biological data into clean categories. This can create the false impression that neuron types are perfectly distinct when, in reality, they might exist on a spectrum. To assess whether the identified neuron groups reflect clearly-defined projection motifs or ambiguous structure, HBMAP returns not only an optimal clustering but also a distribution over plausible groupings, reflecting the possibility of merging, splitting, or reallocating neurons across groups. This uncertainty is summarized by the posterior similarity matrix (PSM), which measures the probability of co-clustering for two neurons both within and across brains. This identifies groups of neurons that are clustered together with high probability (red blocks in Fig. 5e), such as the clearly separated groups of neurons projecting to PM (motif ID: 18) and AL (motif ID: 14) only, as well as uncertainty in further splitting or merging some groups of neurons (yellow areas in Fig. 5e), both within clusters (e.g. the two (LM, AL) motifs, 2 and 15) and across clusters (e.g. yellow areas in the off-diagonal block corresponding to reallocating some neurons across the two (LM, AL) motifs). This insight highlights neurons at the boundary between motifs and helps avoid overinterpretation of weakly supported motifs.

For the optimal clustering, confidence in a neuron’s assignment is measured by the posterior probability of the allocation (Fig. 5f). High confidence is observed for most neurons (high probability in light blue in Fig. 5f). For example, in the AL motif, 75%, 93%, 75%, 93% of neurons for each brain, respectively, have allocation probability greater than 0.9, further supporting the presence of this motif (which was not reported in [11]). Some neurons show lower motif assignment confidence (dark blue in Fig. 5f). This is especially true for the two (LM, AL) sub-motifs where cells with similar projection counts to both regions are harder to classify (Fig. 5f and Supplementary Fig. 3). This suggests that while the data supports distinct sub-motifs based on relative projection strength, the boundary is fuzzy, possibly reflecting true biological overlap or dissection-related ambiguity due to LM and AL’s anatomical proximity.

#### HBMAP enables comparison between barcode sequencing platforms

Recently, combinations of MAPseq and its derivative BARseq enable simultaneous projectome and transcriptome analysis, but since BARseq uses in situ sequencing to identify barcodes while MAPseq relies on extracted RNA, this raises the question of whether results are affected by differences in experimental platform. In [12], projections from the auditory cortex were studied across three brains (one MAPseq, *n*_2_ = 5082 neurons and two BARseq, *n*_1_ = 605 and *n*_3_ = 704 neurons - visualized in Supplementary Fig. 4). HBMAP inferred that with high posterior probability the MAPseq brain (2) exhibits stronger projections to somatosensory cortex and contralateral striatum (SSctx and Cstr) and weaker projections to contralateral auditory cortex and thalamus (AudC and Thal), relative to the BARseq brains (1 and 3) (Fig. 6a). These differences reflect reduced weights for motifs to AudC or Thal only in the MAPseq brain (expected total weight: 0.109 [0.01, 0.12] compared to the BARseq brains, 0.266 [0.21, 0.30] and 0.37 [0.34, 0.41]), and a corresponding increase in the (↑Thal, ↓Cstr) motif suggesting barcode identification method might influence the results. Despite these specific differences, the overall motif abundance profiles remain comparable between methods, as the total variation distances show overlapping CIs (Fig. 6b).

**Fig. 6:**
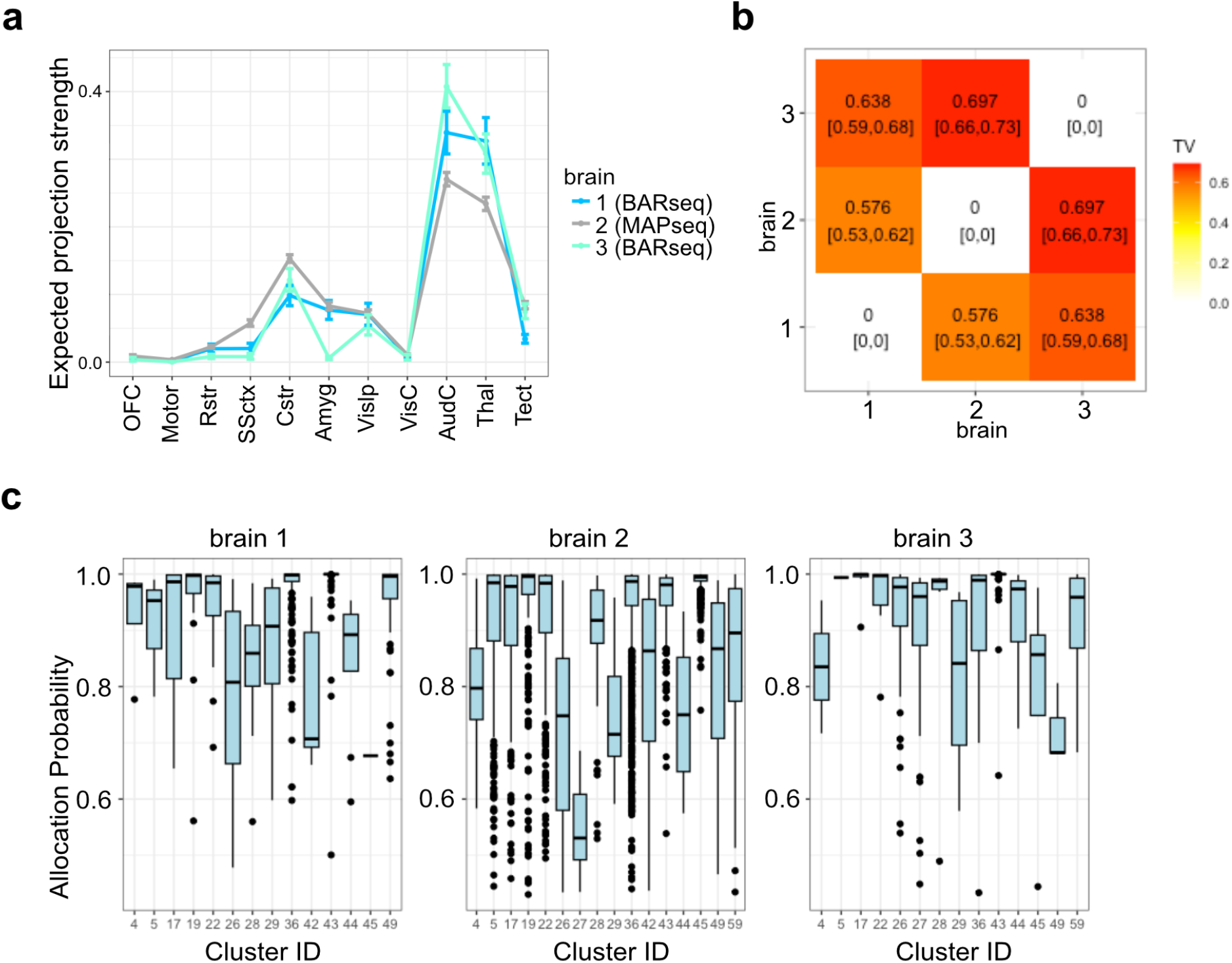
HBMAP enables comparison between MAPseq and BARseq, along with uncertainty in motif assignment. The data [12] considers projections from the auditory cortex to the target regions: the orbitofrontal cortex (OFC), motor cortex (Motor), rostral striatum (Rstr), somatosensory cortex (SSctx), caudal striatum (Cstr), amygdala (Amyg), ipsilateral visual cortex (VisIp), contralateral visual cortex (VisC), contralateral auditory cortex (AudC), thalamus (Thal), and tectum (Tect). **a**. The estimated perregion projection strengths for each brain, along with CIs. **b**. TV distance matrix, illustrating the extent to which the motif structures for different mice diverge. **c**. Boxplots indicating the allocation probabilities of the neurons in each cluster, in each brain. Cluster 43: Thal; 45: ↑Thal,↓Cstr; 44: ↑Thal, ↓Tect; 49: Thal, Tect; 59: Thal,Tect,↓Cstr; a complete labeling and visualization of all motifs is given in Fig. 5.

In the same dataset, HBMAP identifies 63 distinct clusters and 16 prominent motifs, echoing and extending findings from [12] (Supplementary Fig. 5). Using a hierarchical k-means approach informed by known cortical neuron types (corticothalamic (CT), pyramidal tract-like (PT-l) and intratelencephalic (IT)), [12] revealed 25 subclusters, which align well with HBMAP’s motifs with some discrepancies (further description in Supplementary Fig. 5).

The authors assessed the uncertainty in the allocation of neurons into these clusters with a two-stage approach, which first clusters and then fits a classification model to estimate class probabilities, risking overconfidence (see Supplementary Fig. 6). Indeed, they find that 93% of all neurons were uniquely assigned to a single cluster with high probability> 98%, but given the inherent noise in MAPseq, such high confidence in the presence of subpopulations of neurons with unique projection patterns is too optimistic. Instead, HBMAP finds greater uncertainty in the allocation probabilities, which is illustrated for prominent motifs in Fig. 6c and Supplementary Fig. 7. For example, focusing on prominent motifs of the CT and PT-l type, HBMAP shows high confidence in motif allocation among neurons in the CT motifs: the percentage of neurons having a probability greater than 0.8 is 97% overall for the (Thal) motif and 99% overall for the (↑Thal, ↓Cstr) motif. Instead, uncertainty is greater for the motifs related to PT-l; the percentage of neurons having a probability greater than 0.8 is 63% overall for the (↑Thal, ↓Tect) motif, 83% overall for the (Thal, Tect) motif, and 70% overall for the (Thal, Tect,↓Cstr) motif. Therefore, HBMAP confirms diversity in neurons’ projection patterns, but assigns greater uncertainty in specific motifs than previously thought.

## Discussion

We introduced HBMAP, a hierarchical Bayesian model that analyzes high-throughput projectomic data by directly modeling barcode counts to simultaneously infer projection motifs, account for experimental variability, and quantify uncertainty. Applied to MAPseq data from the auditory and visual cortex, HBMAP discovered prominent projection motifs of V1 neurons projecting to a single target, as well as bi-and trifurcating patterns with varying strengths, confirming and extending previous findings. In the auditory cortex, it confirmed projection diversity but revealed greater uncertainty in specific motifs than previously recognized. Unlike existing methods that pool data, HBMAP’s hierarchical structure distinguishes biological signals from experimental variation. Its interpretable and generative design supports both biological insight and experimental planning, including model fit evaluation and synthetic data generation.

HBMAP could be further improved by including information from barcode counts in the injection site, which may provide additional information on the noise level and confidence in projection strength. Another aspect is that projection motifs reflect projection strength relative to the considered target regions. Including more target regions may alter the identified motifs; in particular, target regions with very high barcode counts may obscure differences in regions with smaller counts. Probabilistic multi-view clustering [24] could address this by grouping target regions into views, to elucidate differences within views.

As sequencing-based methods for projection analysis evolve, HBMAP could be further developed to account for multiple injection sites and transcriptomic profiles of barcoded neurons. Moreover, the generative capacity of HBMAP is useful to build tools for sample size determination in novel experiments. The algorithm underlying HBMAP may be computationally demanding if the number of neurons observed is in the hundreds of thousands, and in this case, faster, yet approximate algorithms may be preferred for scalability [25]. Lastly, we note that the Dirichlet-multinomial model has been used in a variety of domains, including genomics [26], language modelling [27], documentation classification and topic modelling [28, 29], and stylometry [30], and mixtures of Dirichlet-multinomials have been applied, for example, to microbial metagenomics [31], text data [32], and solar radiation classification [33]. Therefore, the hierarchical implementations that we develop for HBMAP have potential relevance to extend analysis toolkits in diverse additional domains.

## Methods

HBMAP assumes neuron barcode counts are generated from a multinomial model given neuron-specific projection strengths, which are unknown and modeled with a mixture of Dirichlets:

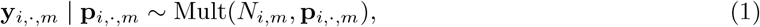

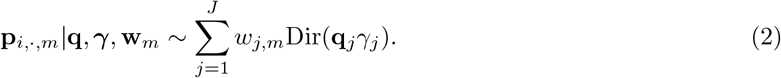

Equivalently, this assumes barcode counts follow a hierarchical mixture of Dirichlet-multinomials:

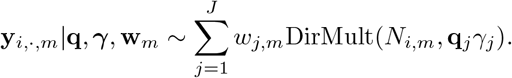

The HBMAP mixture model in (1)-(2) can be augmented with latent allocation variables *z*_*i,m*_ for each neuron *i* and mouse *m*, where *z*_*i,m*_ = *j* if neuron *i* in mouse *m* is generated from the *j*th motif; this provides a way to generate data from the model, by first probabilistically selecting a motif, then simulating the neuron’s projection strength given the mean projection strength and dispersion characterizing the selected motif, and lastly, generating neuron counts given the projection strength;

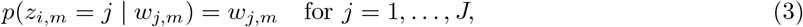

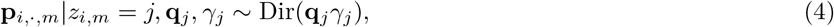

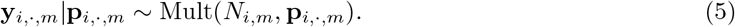

The latent variables 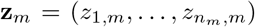characterize a clustering of the neurons into groups for each brain, where clusters may be shared across brains. In model-based clustering, the *kernel* of the mixture model (i.e. the Dirichlet-multinomial model in our case) plays a critical role in obtaining clusters of practical relevance, and therefore, the kernel should be selected carefully to reflect the shape and properties of a cluster for the application at hand. HBMAP employs a Dirichlet-multinomial kernel, which is an intuitive model for MAPseq, as motivated the overview sectin (Fig. 2 and Fig. 3); compared with a multinomial kernel, it allows for the more realistic characterization of a cluster as a group of neurons with similar (but not necessarily the same) projection strengths.

Each mixture component represents a projection motif, with parameters **q**_*j*_ reflecting the mean projection strength within motif and *γ*_*j*_ controlling variability across neurons (as illustrated in Fig. 3e). To discourage the presence of motifs with only small differences in their mean projection strengths, HBMAP employs a repulsive Dirichlet prior on **q**_*j*_:

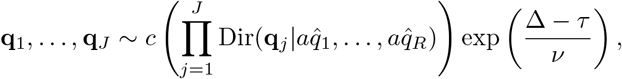

where *c* is the normalizing constant; 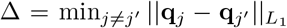is the minimum *L*_1_ distance between the mean projection strengths of all pairs of motifs; and the parameters *τ* and *v* control the repulsion, with larger *τ* encouraging a larger minimum distance and smaller *v* increasing the strength of the repulsion term. The default HBMAP values are *τ* = 0.4 and *v* = 1*/*20, which, for example, promotes that a unicasting motif with strength *q*_*j,r*_ = 1 to region *r* and a bicasting motif with strength 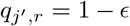to the same region *r* are separated by at least a difference of *ϵ* = 0.2 in the projection strength. Additional parameters of the repulsive Dirichlet prior include the prior mean 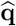, which is empirically set as the average empirical projection strength across all neurons, and the parameter *a* reflecting sparsity (with smaller values encouraging sparse, e.g. unicasting and bicasting, motifs and default value of *a* = 2). A truncated Gamma prior is employed for *γ*_*j*_:

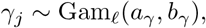

with shape *a*_*γ*_, rate *b*_*γ*_, and lower bound *l*. Recalling that small *γ*_*j*_ implies high variability, the lower bound can be useful to reduce highly overdispersed motifs (default of *l* = 1). The default values of *a*_*γ*_ = Median(*N*_*i,m*_) and *b*_*γ*_ = 2 reflect less overdispersion when overall total barcode counts are higher.

HBMAP allows the abundance of each motif *w*_*j,m*_ to vary across brains, with the hierarchical model allowing for data-driven borrowing of information across brains:

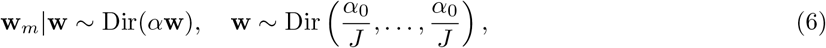

where *J* represents a specified upper bound on the number of motifs. In our experiments *J* is chosen generously based on a simple elbow plot (total within-cluster variation against the number of clusters) of an initial k-means fit; this helps to choose an upper bound *J* which is large enough, but not too large for computationally efficiency. Due to its importance in determining how much information to borrow across brains, the parameter *α* is also learned through the Bayesian framework, with *α ~* Gam(*a*_*α*_, *b*_*α*_). Similarly, *α*_0_, which influences the number of observed motifs, is assigned the prior 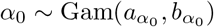. The parameters of the prior on *α* and *α*_0_ can be specified by viewing the sparse overfitted mixture, specified by equations (1)-(6), as a finite-dimensional approximation to the nonparametric mixture model employing a hierarchical nonparametric prior, namely, the hierarchical Dirichlet process [HDP, 34]. Specifically, we make use of the property of the HDP, which states that asymptotically, the number of motifs in each brain (denoted *k*_*m*_) has prior expectation 𝔼 [*k*_*m*_] *≈ α* log(*n*_*m*_) and the number of unique motifs across all brains (denoted *k*) has prior expectation 𝔼 [*k*] *≈ α*_0_ log(*n*_1_ + … + *n*_*M*_) [35]. Based on these equations, initial estimates for the number of motifs both within and across brains can be used to set the prior mean of *α* and *α*_0_.

### Inference Algorithm

A summary of the parameters of HBMAP and their relations is provided in the graphical model (Fig. 3f), where circles with white and colored fills denote global and local parameters, respectively. As exact computation of the posterior distribution of all parameters in HBMAP is not feasible, a Markov chain Monte Carlo (MCMC) algorithm is employed. MCMC is considered to be the gold-standard tool for Bayesian inference, due to its guarantees of producing asymptotically exact draws from the posterior distribution when the algorithm is run long enough. The MCMC algorithm is a Metropolis-within-Gibbs scheme that iteratively cycles through the parameters in blocks, sampling from their full conditional distributions. Specifically, in the graphical model of HBMAP (Fig. 3f), the algorithm iterates through each circled parameter, inferring it based on both the current values of the variables to which it points and the variables which point to it. Full details of the algorithm are given in Supplementary Materials. The MCMC algorithm produces a large number of samples from the posterior; in the following, let the superscript (*t*) of any parameter, e.g. **q**^(*t*)^, indicate the MCMC samples for *t* = 1, …, *T*.

### Clustering and its uncertainty

An advantage of the Bayesian clustering framework [36] is that it provides a posterior over the partition space, i.e. a range of plausible groupings, along with how confident we are in each possibility. In particular, the MCMC algorithm produces an ensemble of clustering solutions, representing draws from this posterior. To obtain a single optimal clustering solution, we minimize the posterior expected variation of information (VI) [37–39], approximated by the MCMC draws (this optimal solution for the visual cortex data is illustrated in Fig. 5e-f). Due to the well-known label-switching issue in mixtures [40], after obtaining this optimal clustering, we run a subsequent MCMC algorithm with the clustering fixed to this solution, in order to investigate the projection motifs and differences across brains.

To quantify confidence in the neuron’s motif assignment, we compute the posterior probability of the neuron’s assignment, given the allocation of all others:

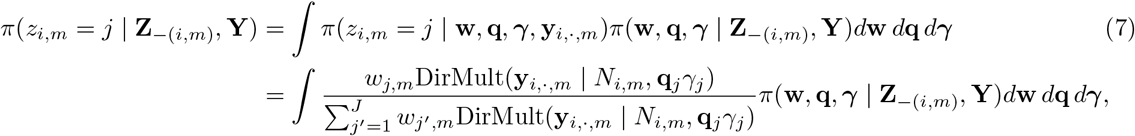

where **Z** and **Y** denote the collection of allocation variables and barcode counts across all brains and the subscript *−*(*i, m*) denotes this collection with the variable for the *i*th neuron in the *m*th brain removed. Assuming the posterior of the parameters is relatively unchanged when the allocation of *i*th neuron is removed, i.e. *π*(**w, q, *γ*** | **Z**_*−*(*i,m*)_, **Y**) *≈ π*(**w, q, *γ*** | **Z, Y**), the posterior allocation probability in (7) is approximated from the MCMC samples by:

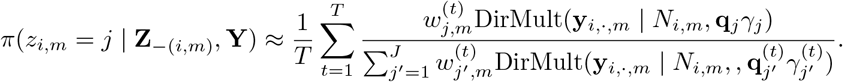

To characterize uncertainty in the clustering structure across the ensemble of posterior samples, we compute a Monte Carlo approximation of the posterior similarity matrix (PSM), with elements representing the co-clustering probability between neurons:

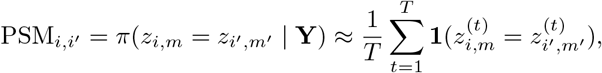

for neuron *i* in brain *m* and neuron *i*′ in brain *m*′,

### Probabilistically identifying prominent motifs

Given the optimal clustering, HBMAP probabilistically identifies prominent and reliable motifs that would be expected to be observed if data were collected for a new brain. This involves computing the posterior probability that the global weight of each motif is greater than a threshold *ϵ*:

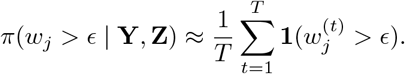

If this probability exceeds a specified value, e.g. 0.95, then motif is considered prominent. Fig. 5b compares this probability with the posterior expected global abundance 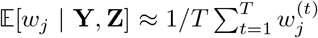for the visual cortex data with *ϵ* = 0.02.

### Projection motif analysis

To understand the motif projection patterns, HBMAP provides a posterior distribution of the mean projection strength **q**_*j*_ for each motif identified in the optimal clustering. Importantly, this provides robust projection motif analysis that accounts for noise and uncertainty due to total barcode counts, compared with simply computing the average empirical projection strength within motif. Specifically, the mean projection strength **q**_*j*_ is estimated, along with CIs, from the MCMC samples 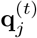as 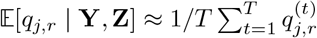.

To reliably determine the projecting regions of neurons within a motif, HBMAP employs a probabilistic rule. First, recall that if

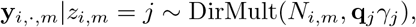

then from compound formulation of the Dirichlet-multinomial,

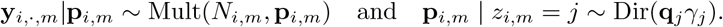

For each motif *j* and target region *r*, HBMAP computes the posterior predictive probability that the projection strength of any additional neuron is greater than a threshold *ϵ*:

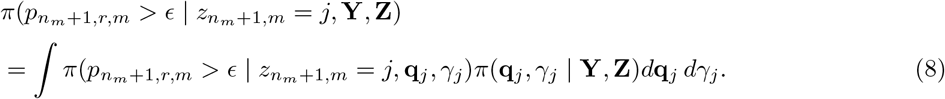

This reflects the idea that if data were collected for any additional neuron, it would project to the identified regions of the motif with sufficiently high probability. The integral in (8) can be approximated based on the MCMC draws, and the integrand is:

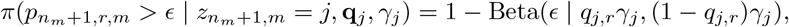

where Beta(*ϵ* | *a, b*) denotes the Beta cumulative distribution function evaluated at *ϵ* with parameters *a, b*. To label motifs with the number and name of target regions, these posterior probabilities in (8) are thresholded at 0.5, that is, region *r* is a target region of motif *j* if

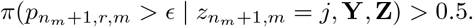

### Quantifying variable motifs and brain differences

#### Variable motifs

To identify variable motifs, that is, motifs whose abundance *w*_*j,m*_ varies across brains, HBMAP computes the posterior expected variance of *w*_*j,m*_ across brains. For each motif identified in the optimal clustering, this is defined as

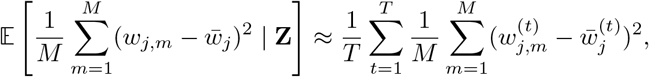

Where 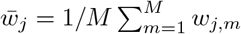. This variance is compared to a null setting, which assumes no brain differences and that the neurons’ motif assignments are generated identically across brains from a simple Categorical model based on the global motif abundance. Specifically, null motif allocations are generated assuming 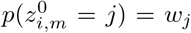. HBMAP computes the probability that expected variance of the brain-specific weights given the optimal clustering is larger than the expected variance of the brain-specific weights given the null motif allocations:

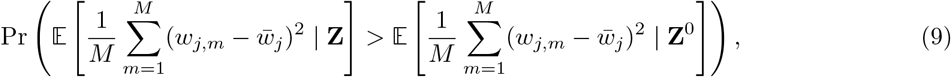

where the probability is with respect to the null model. Here, **Z**^0^ collects the motif assignments across brains and neurons under the null, and (9) is approximated using Monte Carlo by generating multiple motif allocations from the null model. This identifies motifs that are variable (and more stable) across brains (see Fig. 5c which compares the variance of *w*_*j,m*_ across brains to the probability that this variance is large under the null.

#### Quantifying brain differences

To quantify the differences between any pair of brains across all motifs, HBMAP computes the total variation (TV) distance between the brain-specific weights **w**_*m*_. The TV ranges from zero to one, with zero corresponding to equivalence and one if there is no overlap in the projection motifs present in the two brains. It is defined as:

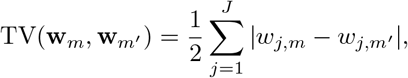

for any two brains *m* = 1, …, *M* and *m*′ = 1, …, *M*. Through the Bayesian framework, we can infer the posterior distribution over the TV distance between any pair of brains and summarize with the posterior expected TV distance, i.e. 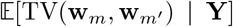, along with CIs. Fig. 5d visualizes the TV distance to quantify experimental differences in the visual cortex data.

#### Brain-specific projection inference

HBMAP also allows for robust inference of brain-specific quantities, such as the posterior expectation or covariance of the projection strength, which summarizes projections across all neurons in the brain. Unlike simply computing the mean and covariance of the empirical projection strength across all neurons in the brain, which ignores confidence and information given by the total barcode counts, HBMAP provides more reliable estimates with sound uncertainty. For example, the posterior predictive expected projection strength to region *r* for brain *m* is:

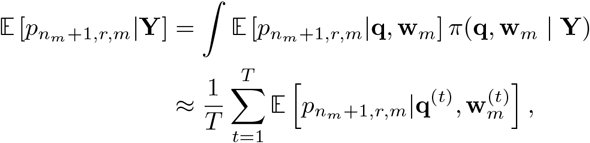

where given (**q, w**_*m*_), we have

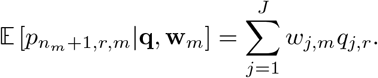

Fig. 6a compares the expected projection strength across brains for the data from [12], finding that the MAPseq brain (2) has overall lower projection strength to AudC and Thal with high probability (non-overlapping CIs).

### Synthetic data generation and posterior predictive checks

The generative nature of HBMAP permits the simulation of synthetic data, denoted **Y**^*s*^, drawn from the posterior predictive distribution:

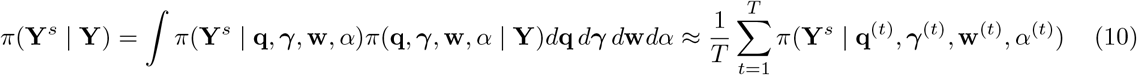

Specifically, for a desired number of brains *M* and neurons *n*_*m*_ in each brain *m* = 1, …, *M*, synthetic data is obtained by first drawing the global parameters from the posterior distribution, which is achieved by randomly choosing the parameters **q** = **q**^(*t*)^, ***γ*** = ***γ***^(*t*)^, **w** = **w**^(*t*)^, *α* = *α*^(*t*)^ of an MCMC sample, and then generating the local brain-specific data from the model given those parameters following equations in (3) - (6), namely, the brain-specific motif abundances, allocations, projection strengths, and counts for the synthetic data are:

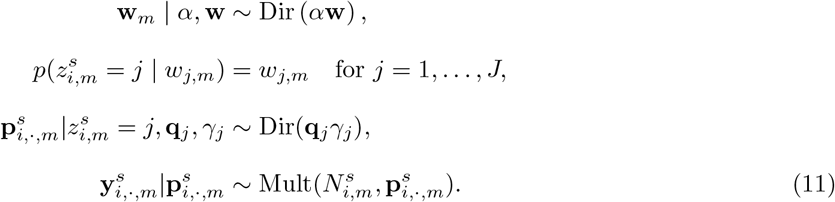

Note that generating synthetic barcode counts from the multinomial in the last step above (11) requires having the neuron’s total barcode count. Thus, each neuron’s total barcode count is also simulated by sampling from a truncated negative binomial,

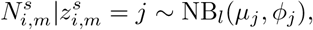

where the cluster-specific mean *µ*_*j*_ and dispersion *ϕ*_*j*_ are estimated via a method of moments. The lower bound *l* reflects the minimum total barcode count used when preprocessing the data (if applied).

These synthetic data are useful for posterior predictive checks to assess the fit of HBMAP to the observed data. Specifically, the empirical projection strengths of the neurons, computed as 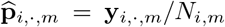, are compared between the observed and synthetic data (for example, through boxplots and histograms in Supplementary Fig. 3); similarity supports HBMAP, while differences highlight potential limitations in the model. Additionally, the synthetic data can be used to test hypotheses and robustness (Fig. 4c,d) or for sample size calculations.

### Preprocessing of MAPseq

We perform the same preprocessing steps for the two datasets describing projections from the visual cortex [11] and auditory cortex [12] as carried out in the respective papers.

For the data from [11], the following filtration steps are performed on the raw barcode count data:

- Apply spike-in normalization to the raw barcode count matrices.
- Remove any neuron without a barcode count of at least 10 in at least 1 target site.
- Remove any neuron without a barcode count of at least 300 at the injection site (V1).
- Remove any neuron with less than ten-fold difference in barcode counts between V1 and the most barcode rich target site.
- Subset the raw barcode count matrix to only include neurons which obey the above criteria, and then normalize relative to the spike-in counts observed in the negative control sample (olfactory bulb).

For the data from [12], the following filtration steps are performed on the raw barcode count data:

- Remove any neuron with less than 10 or more than 10,000 barcodes at the most barcode-rich target site.
- Match barcode sequences at the target site to those at the injection site, allowing either 3 mismatches (MAPseq-XC14) or 1 mismatch (BARseq - XC9 and XC28).
- Remove neurons collected from below the intended injection site (BARseq).
- Remove neurons from highly distorted brain slices (BARseq).
- Apply spike-in normalization.

## Data availability

Data is available for both datasets from the respective papers, [11] and [12]. The processed datasets are both included in the accompanying **R** package https://github.com/huizizhang949/HBMAP.

## Code availability

The source code of the HBMAP project is available via GitHub at https://github.com/huizizhang949/ HBMAP, where also the implementation of the binomial model and k-means model and detailed deployment instructions are provided.

## Acknowledgements

G.S. is funded by a Wellcome Trust Sir Henry Dale Fellowship 211236/Z/18/Z. S.W. acknowledges funding by the Engineering and Physical Sciences Research Council (EP/Y028783/1). E.A. is funded by a Medical Research Council Precision Medicine DTP Fellowship. We thank Matthew Nolan for insightful feedback on the manuscript.

## Author information

### Authors and Affiliations

**School of Mathematics and the Maxwell Institute, University of Edinburgh, Edinburgh, UK**

Sara Wade, Jinlu Liu, and Huizi Zhang.

**Centre for Discovery Brain Sciences and Simons Initiative for the Developing Brain, University of Edinburgh, Edinburgh, UK**

Gülşen Sürmeli and Edward Agboraw.

**Simons Initiative for the Developing Brain, University of Edinburgh, Edinburgh, United Kingdom**.

Gülşen Sürmeli

## Contributions

S.W. and G.S conceived of the method design and analyses, with input from all authors. S.W., J.L., and

H.Z. wrote the code and accompanying software and performed the computational analyses, except where otherwise noted. E.A. performed data preprocessing and the binomial model implementation and analysis.

S.W. and G.S. wrote the manuscript with input from all authors.

## Supplementary information

Supplementary Notes 1–2 and Figs. 1–7.

## Supplementary Information

### 1 Preprocessing and Filtration

For both datasets, standard filtration steps are carried out to remove certain neurons (e.g. with low counts) and corrected for sample-specific biases via spike-in normalization, following the steps outlined in the original studies.

### 2 Posterior Inference Algorithm

A Markov Chain Monte Carlo algorithm is developed for full posterior inference. The algorithm is a Gibbs sampler which produces asymptotically exact samples from the posterior by iteratively sampling the parameters in blocks. Let **Z** and **Y** denote the collection of allocation variables and barcode counts across all brains. The full conditional distributions are as follows:

- The allocation variables *π*(**Z**|**w**_1:*M*_, **q, *γ*, Y**);
- Unique parameters *π*(**q**|***γ*, Z, Y**) and *π*(***γ***|**q, Z, Y**);
- Dataset-specific component probabilities *π*(**w**_1:*M*_ |**Z**, *α*, **w**);
- Component probabilities *π*(**w**|**w**_1:*M*_, *α*_0_, *α*);
- Concentration parameters *π*(*α*|**w**_1:*M*_, **w**) and *π*(*α*_0_|**w**);

Simulations of concentration parameters and component probabilities (*α*_0_, *α*, **w**_1:*M*_ and **w**) are the same as in the Norm-HDP model [5].

#### 2.1 Allocation Variables

Based on the full conditionals, the allocation probability of neuron *i* in mouse *m* being allocated to component *j* is:

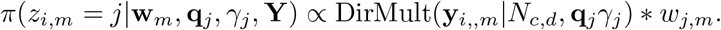

Let

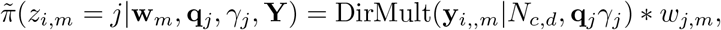

then

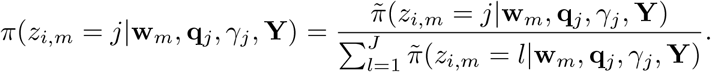

We compute the allocation probability of all *J* components. The posterior distribution of *z*_*i,m*_ follows a Categorical distribution with parameters equal to those *J* values.

#### 2.2 Mean Projection Strengths

Let **q**_*−j*_ denote the projection probabilities for all components except for component *j*. The full conditional distribution for the projection probabilities is:

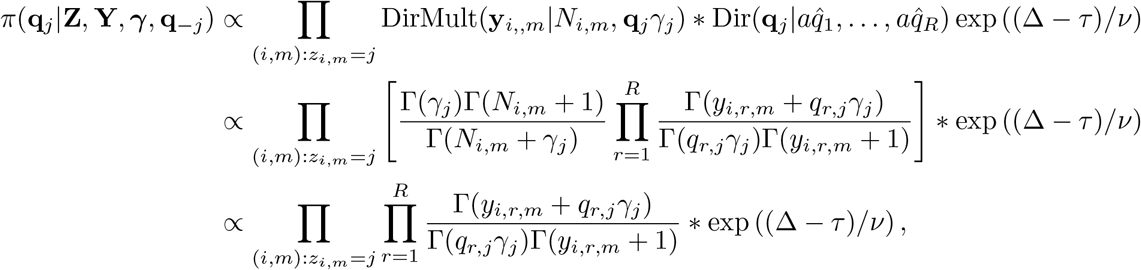

and log of the above full conditional distribution is:

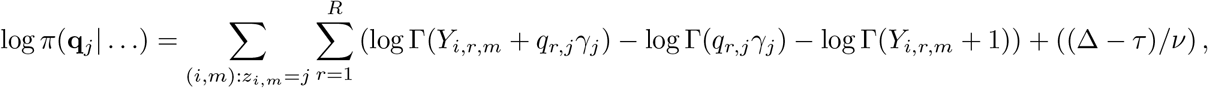

plus some constant. The full conditional distribution for the projection strengths have no closed-form. We apply adaptive Metropolis Hastings to obtain posterior samples. We first transform the unknown parameter **q**_*j*_ into **x**_*j*_, where **x**_*j*_ = (*x*_1,*j*_, …, *x*_*R−*1,*j*_). The transformation involved here is:

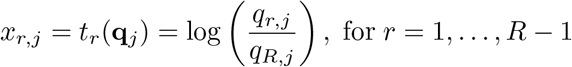

Hence we have:

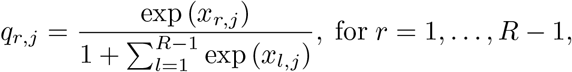

and 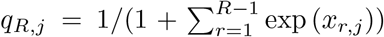. Then a new value for the transformed variable is proposed using a random-walk Metropolis-Hastings where the covariance matrix is updated at each iteration to facilitate convergence of the chain [3, 1]. The computation of acceptance probabilities and updated covariance structure are similar to the posterior inference of the component probabilities in [5].

#### 2.3 Dispersion

The full conditional distribution for *γ*_*j*_ is:

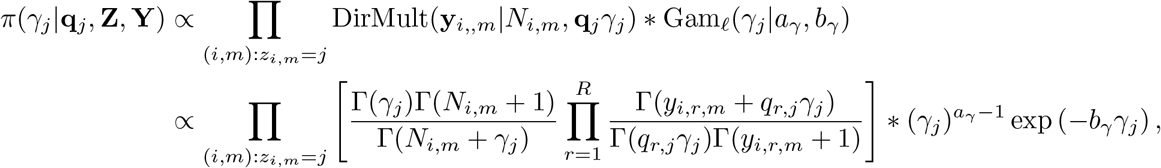

and log of the above full conditional distribution is:

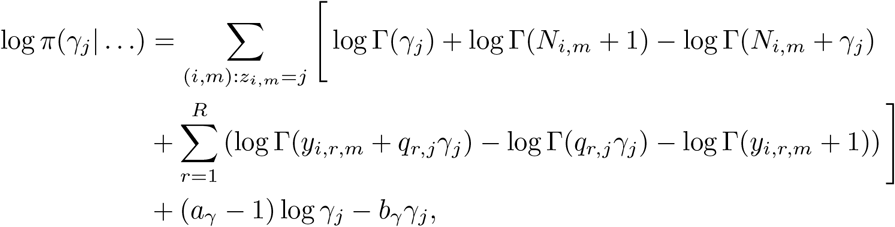

plus some constant. The full conditional distribution does not have a closed-form, and hence adaptive Metropolis-Hastings is applied. A log-transformation is applied to *γ*_*j*_ and the remaining steps are similar to the sampling of the mean projection strengths.

#### 2.4 Dataset-specific Component Probabilities

The full conditional distribution for the dataset-specific component probabilities is:

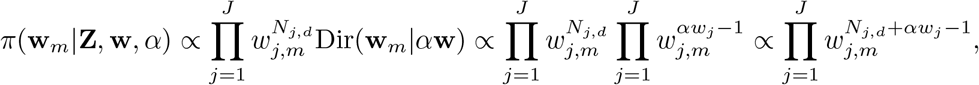

where *N*_*j,d*_ is the number of neurons in component *j* from brain *m*. This is a Dirichlet distribution with parameters equal to *N*_*j,d*_ + *αw*_*j*_, for *j* = 1, …, *J*.

#### 2.5 Component Probabilities

The full conditional distribution for the component probabilities is:

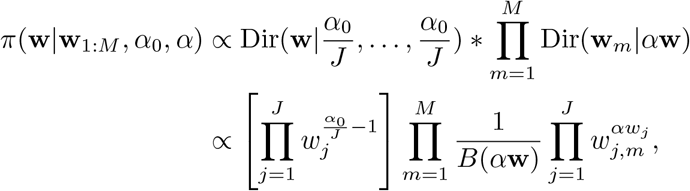

where *B*(*·*) is the multivariate beta function. The log of the full conditional can be written as:

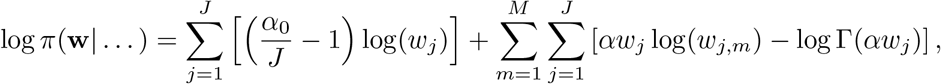

plus some constant. As the full conditional distribution has no closed-form, we will use adaptive Metropolis Hastings to obtain posterior samples of **w**. The transformation applied is the same as the mean projection strengths.

#### 2.6 Concentration Parameters

The full conditional distribution for *α* is:

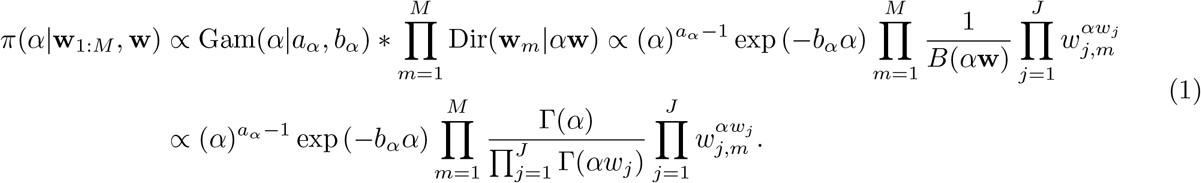

The full conditional distribution for *α*_0_ is:

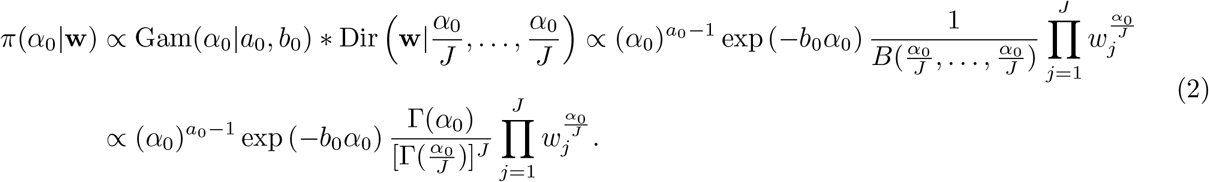

The log of the full condition in eq. (1) is:

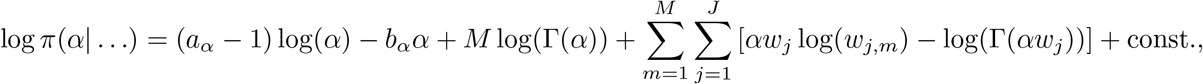

and in eq. (2) is:

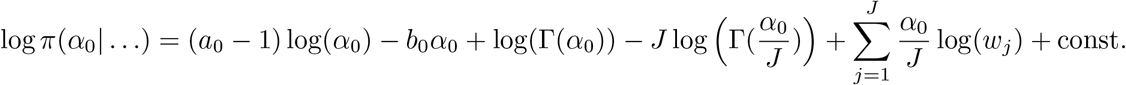

We obtain no closed-form distributions for both concentration parameters *α* and *α*_0_, hence we apply adaptive Metropolis-Hastings to obtain posterior samples. For both parameters, we apply a log transformation.

### Supplementary Figures

**Supplementary Fig. 1:**
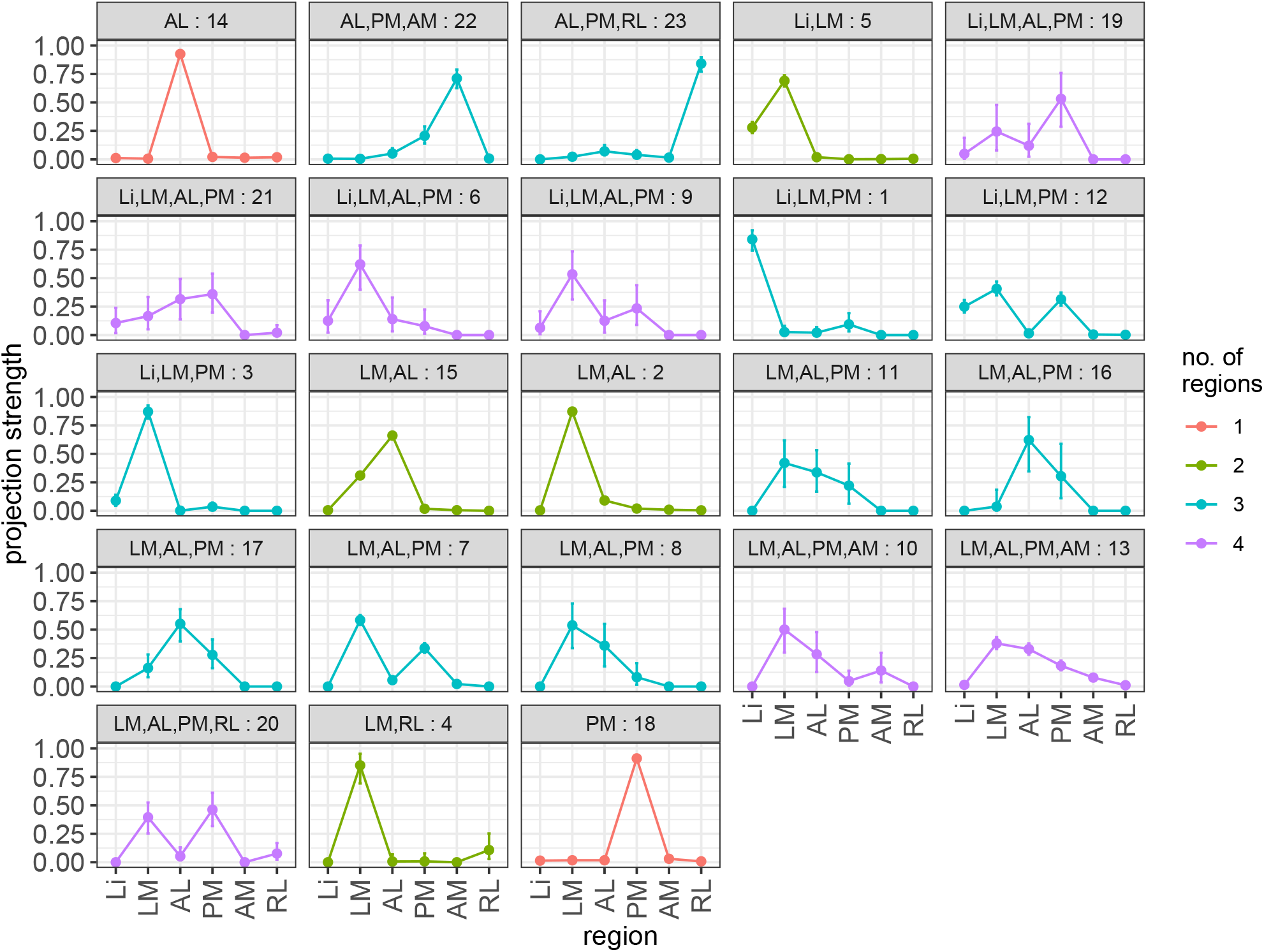
HBMAP probabilistically labels projection motifs. Estimated projection strength of each cluster, along with CIs and colored by the number of projection regions with the identified target regions for each motif provided in each panel title. Results displayed for data from [4].

**Supplementary Fig. 2:**
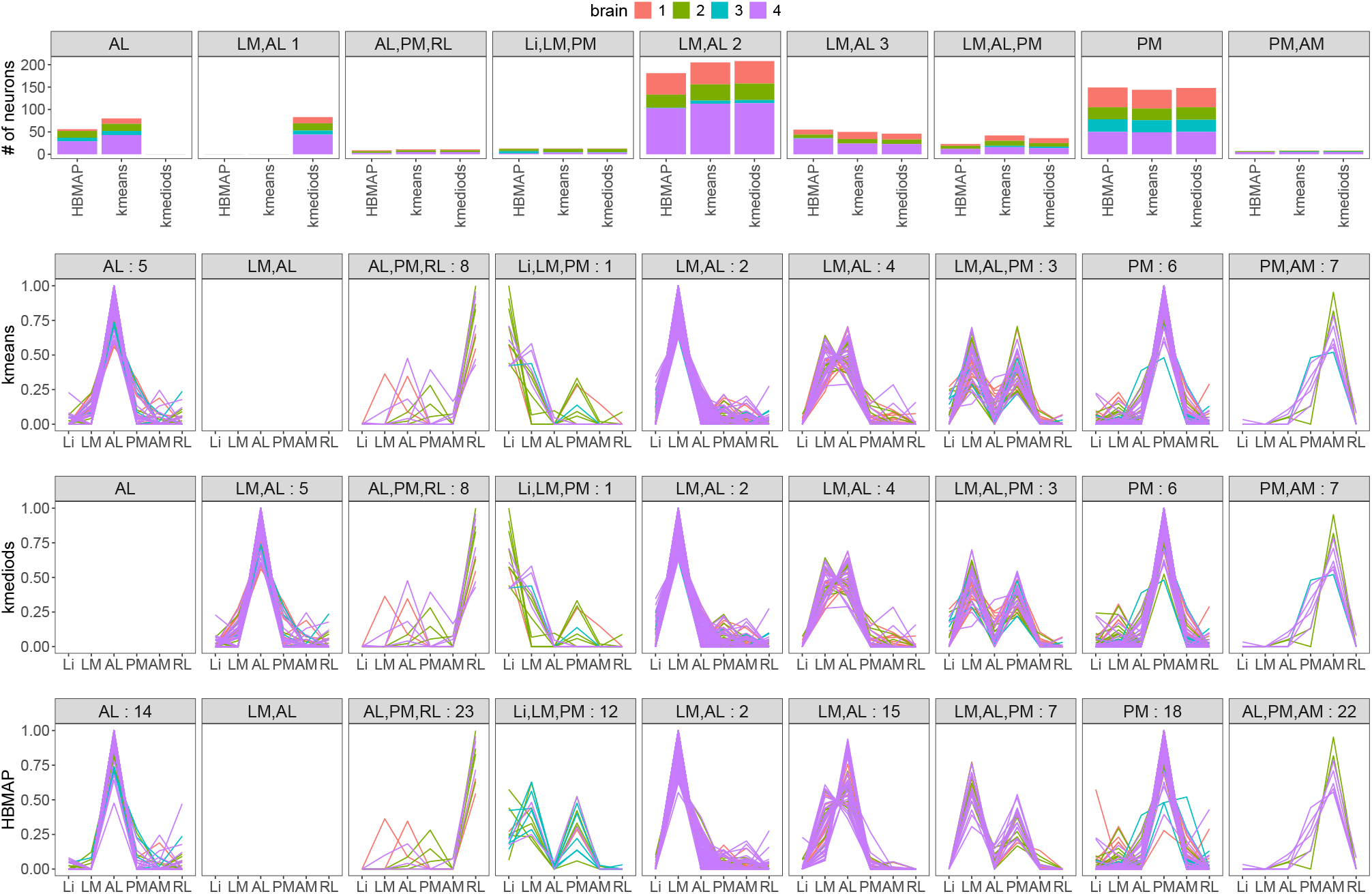
Comparison of the motifs found by k-means, k-medoids, and HBMAP. Results displayed for data from [4]. Top row: number of neurons in each motif across the three methods. Bottom rows: empirical projection strength of neurons within each motif (column) for each model (row). For k-means and k-medoids, the regions of each motif (listed in the panel titles, along with the cluster number) are determined by thresholding the average empirical projection strength across neurons in the motif at 0.05. For HBMAP, the regions of each motif are identified by a probabilistic rule, ensuring that neurons in the motif project to the region with sufficiently high probability.

**Supplementary Fig. 3:**
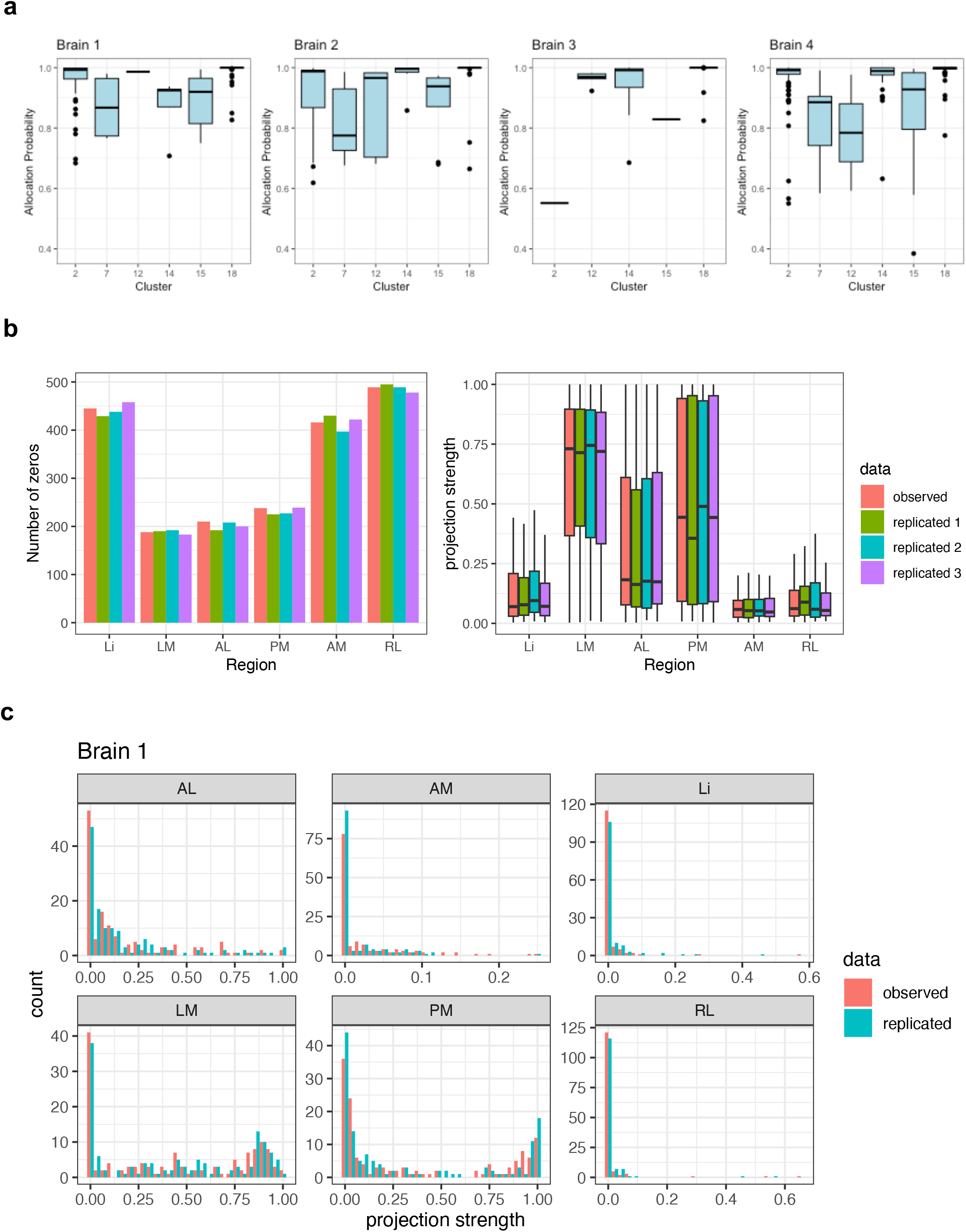
HBMAP provides measures of confidence in motif allocation and tools to assess model fit. Results displayed for data from [4]. **a**. Box-plot of allocation probabilities for the prominent HBMAP motifs across brains; allocations probabilities are greater than 0.5 for almost all neurons. **b-c**. Model fit is assessed through posterior predictive checks, comparing synthetic data generated from HBMAP to the observed data. **b**. Similarity of the distribution of zero and nonzero empirical projection strengths. **c**. Histograms of the empirical projection strengths from a single synthetic dataset.

**Supplementary Fig. 4:**
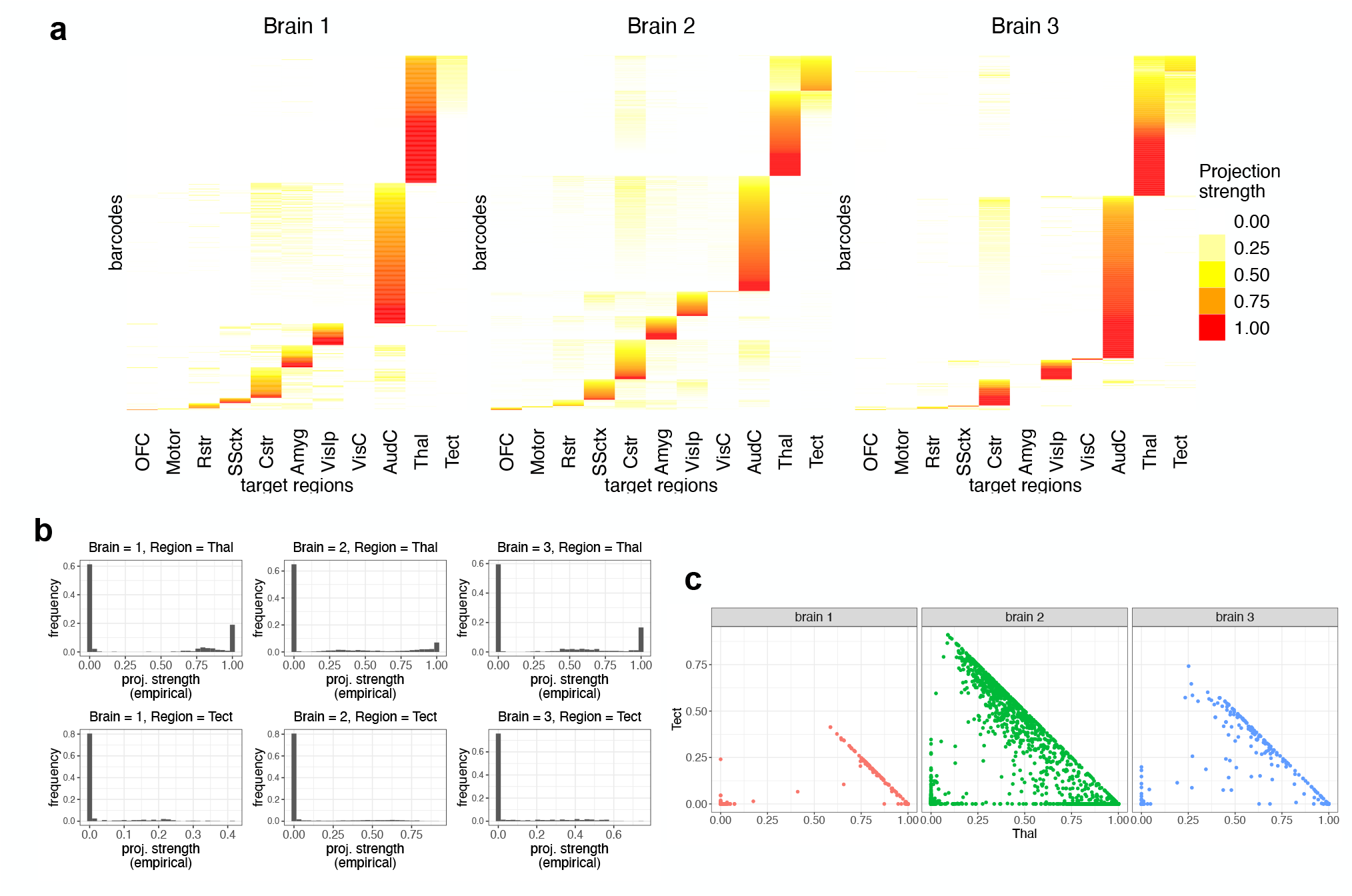
MAPseq data for the auditory cortex displays characteristic features of MAPseq data. **a**. Heatmaps visualizing the empirical projection strengths from normalized barcode counts in brain 1 (BARseq), brain 2 (MAPseq), and brain 3 (BARseq) in the data from [2]. Notable brain-level differences in projection can be observed. **b**. Histograms depicting the empirical projection strengths of neurons in the data from [2] the regions Thal and Tect. Multimodality and overdispersion are evident. **c**. Scatter plot of the empirical projection strengths to two regions Thal and Tect, illustrating sparsity and experimental differences in the data.

**Supplementary Fig. 5:**
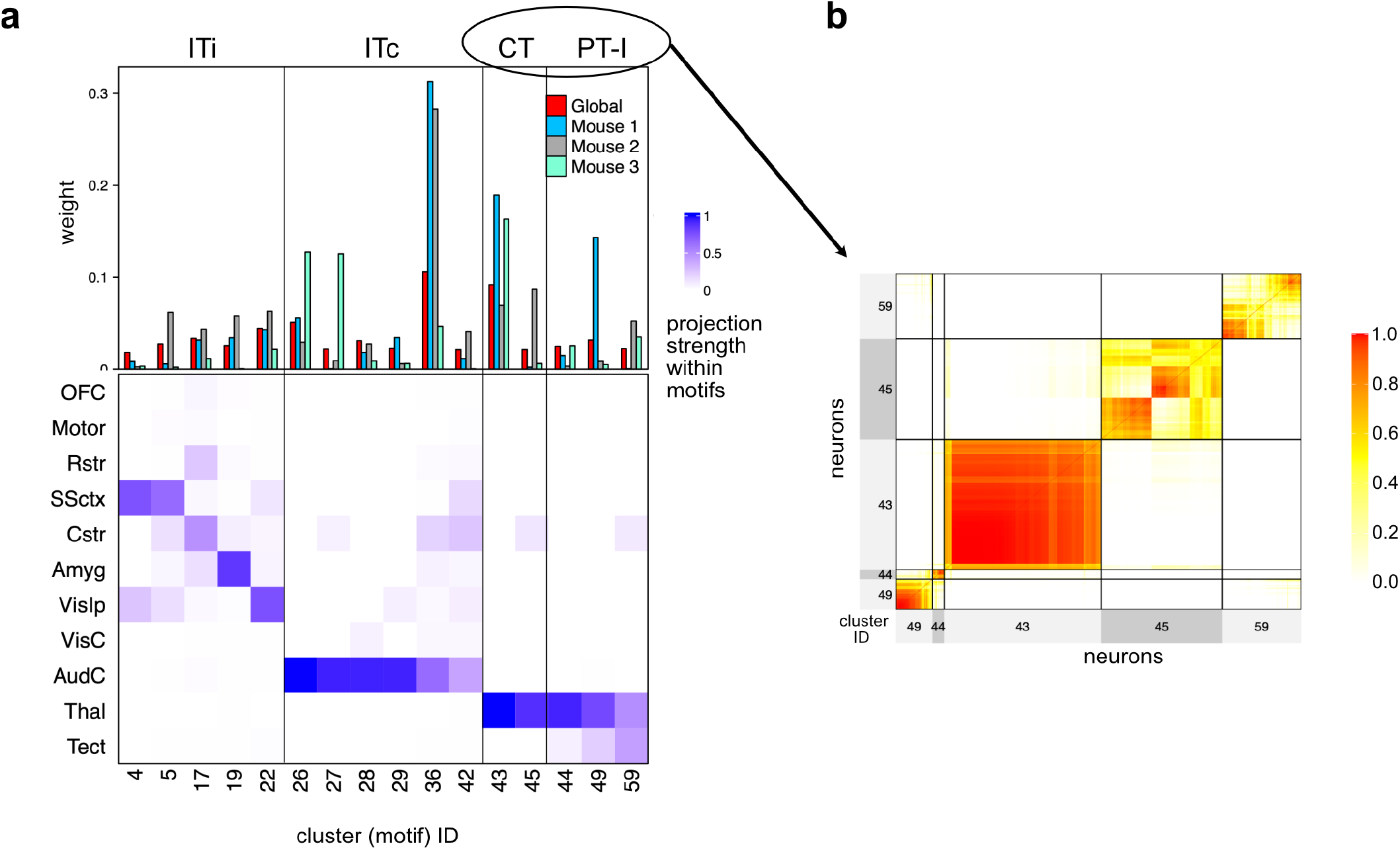
Projection motifs from the auditory cortex align with previously identified classes. **a**. Heat map of the estimated projection strength to the 11 regions for each motif (column), with the estimated global and brain-specific weight attached to each motif visualized above and the estimated brain-specific projection strength to each region on the left. MAPseq is brain 2 and BARseq brains are 1 and 3. The blocks in **a** allocate prominent HBMAP motifs into four major classes, informed by the findings of the original paper [2]. **b**. Posterior similarity matrix for neurons in the CT and PT-l classes. Focusing on the PT-l branch, HBMAP finds three prominent motifs compared to the four motifs of [2] In particular, HBMAP uncovers nuanced variations in the thalamus and tectum (Thal, Tect) motif of [2], splitting it into two submotifs with different modes of projection strength. Moreover, although both approaches identify the PT-l motif to (Thal, Tect, Cstr), there is evident uncertainty in HBMAP’s posterior similarity matrix on further splitting (cluster 59 in PSM on the right), suggesting that this motif may contain neurons with more diverse projections. [2] find two other PT-l motifs, which are also present in HBMAP, but not among the prominent motifs due to very low weights. For the ITc branch, while all motifs in [2] have high and low projection strength to the contralateral auditory and visual cortex (AudC, VisC), respectively, this is not the case for HBMAP, which indeed finds a motif to AudC only. Overall, HBMAP’s findings support and refine the motif categories proposed by [2].

**Supplementary Fig. 6:**
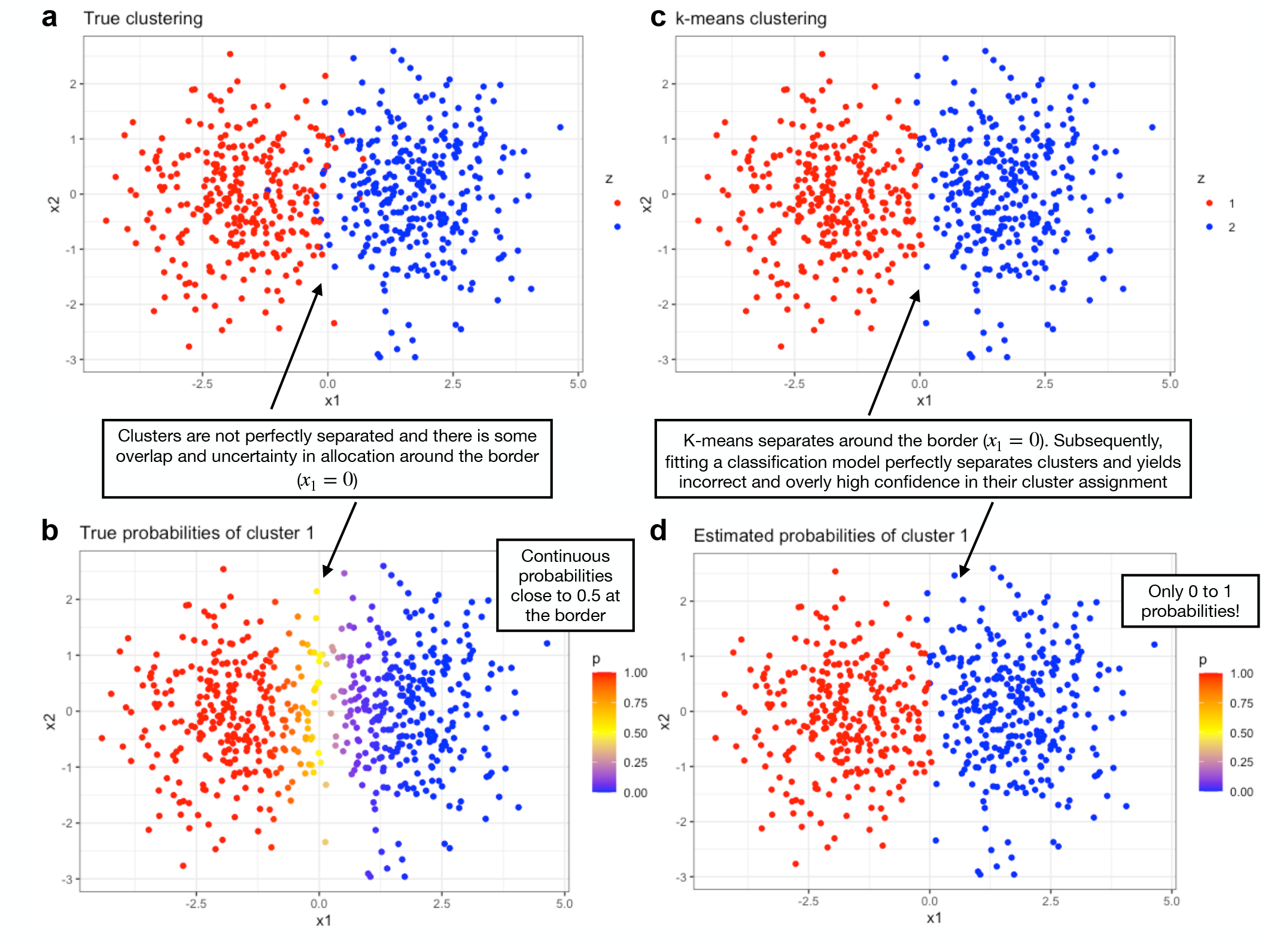
A two-stage clustering and classification approach yields uninformative measures of uncertainty in cluster allocation. While previous literature attempted to assess uncertainty in motif allocation by first clustering and subsequently fitting a classification model to yield class probabilities, this risks optimistic measures of certainty in motif allocations. **a**. As an illustration, data are simulated from two true clusters. **b**. Due to the overlap at the boundary of the clusters, there is some uncertainty in the cluster allocation (probabilities around 0.5), under the true data generating process. **c**. Clustering algorithms separate the space into distinct regions (e.g k-means performs a Voroni tesselation of the space). **d**. Subsquently fitting a classification model confirms that each cluster is associated with a distinct region, yielding probabilities close to 0 or 1, which do not reflect confidence in cluster assignment.

**Supplementary Fig. 7:**
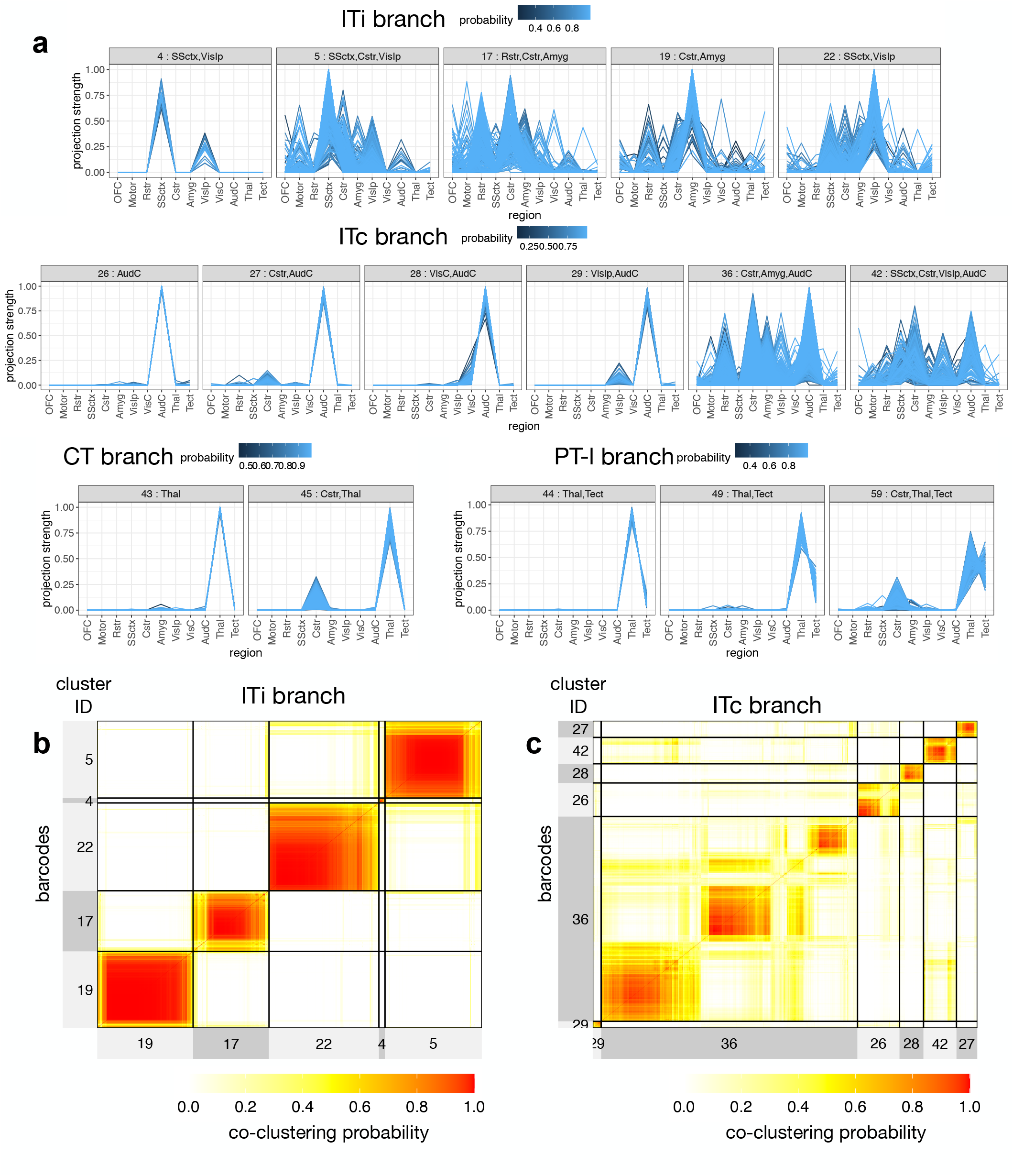
Projection motif assignments from the auditory cortex show greater uncertainty than previously thought. Results displayed for the data from [2]. **a**. Empirical projection strength line plots for the prominent motifs colored by the probability of the neuron’s assignment to the motif. **b-c**. Posterior similarity matrix of neurons in the prominent motifs within the ITi and ITc branches, respectively, representing the probability of co-clustering. For the ITc branch, HBMAP identifies fewer prominent multicasting motifs than [2], but the posterior similarity matrix highlights uncertainty in further splitting them (clusters 36 and 42 in **c**).

